# Resistance of E2F4DN to p38^MAPK^ phosphorylation attenuates DNA damage-induced neuronal death via Cited2

**DOI:** 10.1101/2024.10.29.620791

**Authors:** Aina M. Llabrés-Mas, Noelia López-Sánchez, Alberto Garrido-García, Vanesa Cano-Daganzo, José M. Frade

**Author notes:** Correspondence: José M. Frade.

## Abstract

E2F4 is a transcription factor that supports cellular homeostasis and serves as a substrate for the stress-activated kinase p38^MAPK^, which phosphorylates it at a conserved Thr248/Thr250 motif. A non-phosphorylatable mutant, E2F4DN (Thr248Ala/Thr250Ala), has shown therapeutic efficacy in a murine model of Alzheimer’s disease (AD). We hypothesized that phosphorylation of E2F4 disrupts its protective function, whereas E2F4DN retains this activity during cellular stress. To test this hypothesis, we treated N2a-derived neurons with camptothecin (CPT) to induce genotoxic stress. CPT activated p38^MAPK^ within 8 hours, leading to E2F4 phosphorylation at the Thr248/Thr250 site. We then overexpressed E2F4DN or a phosphomimetic variant, E2F4CA (Thr248Glu/Thr250Glu), and assessed apoptosis by procaspase-3 cleavage. E2F1, used as a pro-apoptotic control, strongly induced caspase-3 activation. This effect was partially mimicked by E2F4CA, whereas E2F4DN markedly suppressed caspase-3 cleavage. Notably, E2F4DN but not E2F4CA upregulated the antiapoptotic factor Cited2, and *Cited2* knockdown abolished the protective effect of E2F4DN. These findings suggest that p38^MAPK^-mediated phosphorylation of E2F4 promotes neuronal apoptosis, while E2F4DN maintains homeostatic function via Cited2, offering mechanistic insight into its neuroprotective role in AD.

## Introduction

E2F4 is a member of the E2F family of transcription factors, which also includes E2F1, a canonical regulator of the G1/S phase transition in proliferative cells with capacity to induce cell death in differentiated neurons [1,2]. E2F4 was initially described as a cell cycle regulator crucial for the establishment of quiescence in proliferative cells [3]. Nevertheless, this transcription factor can regulate more than 7,000 genes [4] that play important roles in cell differentiation, maintenance of DNA integrity, RNA processing, apoptosis, ubiquitination, and cell stress response. Therefore, E2F4 is currently acknowledged as a major player in cell and tissue homeostasis and tissue regeneration [5–8].

There is growing evidence that E2F4 has a multifactorial impact in neuronal welfare and brain homeostasis [9,10], and that this transcription factor may be involved in the physiopathology of neurodegenerative diseases, such as Alzheimer disease (AD), and brain aging [7,8]. E2F4 regulates most of AD-specific gene networks in humans [11], and becomes upregulated in human AD neurons [12] as well as in neurons derived from human induced pluripotent stem cells obtained from familial AD patients [13]. Furthermore, the expression of E2F4 is enriched in blood from MCI and AD [14]. Finally, E2F4 participates in the protective action of memantine and donepezil against cognitive impairment induced by nine common endocrine-disrupting chemicals [15].

The regulation of E2F4 is complex, as it can be post-translationally modified through acetylation and methylation [8]. In addition, E2F4 is susceptible to become phosphorylated by several protein kinases [16–18], and the phosphorylation of E2F4 by the stress kinase p38^MAPK^ within an evolutionary-conserved threonine motif (Thr248/Thr250 in the human sequence and Thr249/Thr251 in mouse) can induce cell cycle re-entry in neurons [16]. This latter effect can be induced by a phosphomimetic form of E2F4 carrying Glu residues in substitution of the conserved threonines (referred to as E2F4CA), and inhibited by a dominant negative form of E2F4 (referred to as E2F4DN) carrying alanines instead of the conserved Thr residues [16].

There is indirect evidence that the Thr conserved motif of E2F4 can be phosphorylated in AD-affected human neurons, based on an E2F4-specific proximity-ligation assay (PLA) [10]. Moreover, a phospho-Thr249-specific antibody generated in our laboratory was able to detect the phosphorylated form of E2F4 in the brain of homozygous 5xFAD mice [9], a known murine model of Alzheimer’s disease (AD) [19] with aggravated phenotype due to the increase of gene dosage [20]. E2F4 phosphorylation at this conserved motif is not specific from the nervous system. Thus, the phosphorylation of Thr248 in human E2F4 and of Thr249 in the mouse sequence has been detected, in retinal pigment epithelium-derived cells and in embryonic stem cells, respectively [18].

We have demonstrated that E2F4DN is an effective and safe multifactorial therapeutic agent in 5xFAD mice. Among the multiple effects of E2F4DN in these mice, we observed reduction of amyloid deposits, gliosis, inflammation, neuronal death, oxidative stress and neuronal tetraploidization, and prevention of dendritic and synaptic alterations, being able to maintain synaptic long-term potentiation [21]. It also extended the life span in both wild-type (WT) and 5xFAD mice, prevented body weight loss and promoted fur regeneration in senile mice [8–10].

Our current hypothesis for the mechanism of action of E2F4DN is that the phosphorylation of the Thr248/Thr250 motif by the stress kinase p38^MAPK^ prevents the homeostatic function of endogenously-expressed E2F4, while the presence of exogenous E2F4DN can restore this function. In this study, we have begun to test this hypothesis by focusing on Neuro2a (N2a) neuroblastoma cells [22] cultured under low serum conditions to induce neuronal differentiation (therefore, we refer to them as N2a neurons). N2a neurons were treated with camptothecin (CPT) to induce DNA damage, as a mechanism of cell stress followed by cell death in neurons [23]. This effect is potentiated by E2F1 expression in a cell cycle-independent manner [2,24,25], while E2F4 can regulate E2F1 function in this cellular stress paradigm [26]. We found that, under these conditions, p38^MAPK^ activity becomes activated in N2a neurons, and this activation correlates with the phosphorylation of endogenous E2F4 at Thr249, an effect that is prevented by the p38^MAPK^-specific inhibitor SB203580. The significance of E2F4 phosphorylation in apoptosis was tested by overexpressing either the phosphomimetic form of E2F4 (E2F4CA) or E2F4DN, followed by quantification of procaspase-3 cleavage as a readout of apoptosis. E2F1, which was used as a positive control, induced a statistically-significant increase of procaspase-3 cleavage 48 h after CPT treatment. This was partially replicated by E2F4CA, while E2F4DN expression inhibited this effect. Finally, we provide evidence that Cbp/P300-interacting transactivator with Glu/Asp-rich carboxy-terminal domain 2 (Cited2) is required for the antiapoptotic effect of E2F4DN in this cellular paradigm. We conclude that the phosphorylation of E2F4 prevents its Cited2-dependent survival effect in N2a neurons subjected to genotoxic stimuli, while E2F4DN rescue these cells from apoptosis. This provides support to the hypothesis that neuronal expression of E2F4DN maintains the homeostatic function of E2F4 under stress conditions.

## Materials and Methods

### Antibodies

Primary and secondary antibodies together with detailed information about their manufacturers and dilutions that were used for the different procedures in which they were used are shown in Supplementary Tables 1 and 2, respectively. The immunopurified rabbit polyclonal antibody recognizing the phosphoThr249 epitope of mouse E2F4 (p-E2F4) was previously described by [9].

### Peptides

A control peptide containing the transactivator of transcription (TAT) sequence of human immunodeficiency virus-1 (HIV-1) (TAT peptide: YGRKKRRQRRR), and a peptide containg the TAT sequence followed by the amino acids 237-256 of the mouse E2F4 sequence (NCBI accession number NP_683754) containing the Thr249Ala/Thr251Ala mutation (TAT-E2F4DN peptide: YGRKKRRQRRRQESSPPSSPQLTAPAPVLGS) were produced by Synpeptide Co. Ltd.

### Adenoviral vectors

The coding sequence of human E2F1 (NCBI accession number NM_005225), preceded by the HA tag, or of human E2F4 (NCBI accession number NM_001950), followed by the 6xHis taq, were synthesized by GenScript and cloned into the pcDNA3 plasmid (Invitrogen) to generate the pE2F1 and pE2F4 plasmids. Then, two mutations were introduced into the pE2F4 plasmid to generate the pE2F4DN (Thr248Ala/Thr250Ala) and pE2F4CA (Thr248Glu/Thr250Glu) plasmids. Finally, serotype 5 adenoviral vectors (Ad5) carrying the human E2F1, wild-type E2F4 (E2F4WT), E2F4DN, and E2F4CA coding sequences were generated as previously described by [27]. To this aim, recombinant plasmids were generated for the coding sequence of human each of these proteins flanked by Adenovirus-specific inverted terminal repeats (ITRs) as well as all the genes necessary to generate the vector, except for E1, which codes for the E1A and E1B proteins. These latter proteins were constitutively expressed by the Q packaging cell line BI-HEK_293A, which was transfected with the respective recombinant plasmids described above. In addition, an Ad5 vector expressing a mouse *Cited2*-specific shRNA carrying EGFP as a reporter (Ad5-m-CITED2-shRNA; Cat. No: shADV-255523) was purchased to Vector Biolabs.

### Cell cultures

Mouse N2a cells [22] were maintained in Dulbecco’s modified Eagle medium (DMEM) (Gibco) supplemented with 10 % fetal bovine serum (FBS) (Gibco), and 0.5% penicillin/streptomycin (P/S) (Gibco). A maximum of 15 passages were given to each cell batch to maintain the characteristics of the cells. For neuronal differentiation, N2a cells were plated at 20,000 cells/cm^2^ in DMEM supplemented with 1% FBS without antibiotics and cultured for 18-20 h before performing the experiments. For immunocytochemistry, cells were plated at the same conditions as above on coverslips (Menzel Glässer, 10 mm) pretreated with 65% nitric acid for 18-20 h, and then extensively washed with MilliQ water and sterilized with absolute ethanol.

### Adenoviral transduction

For transduction, N2a neurons were cultured for 48 h in the presence of 85 infectious units (IU)/cell of each Ad5 vector. For co-transduction experiments, 60 IU/cell were used for each Ad5 vector (120 IU in total). Around 75-95% of cells were efficiently transduced with these infectious titers. Biosecurity procedures have been approved by the Ethics Committee of CSIC.

### DNA damage induction

To induce severe DNA damage to N2a neurons, 10 µM CPT (Cell Signaling Technology), prepared in dimethyl sulfoxide (DMSO), was added to the cultures during different time points depending on the experimental design. CPT is a known topoisomerase I inhibitor [28] that causes double-strand breaks in the replication forks that are associated with DNA replication as well as in the DNA supercoils that are formed during transcription [29].

### p38^MAPK^, c-Jun N-terminal kinase (JNK), and proteasome inhibition

The p38^MAPK^ inhibitor SB203580 (Cell Signaling Technology) was used at 10 µg/µl, and the JNK kinase inhibitor SP600125 (Calbiochem) was used at 25 µg/µl. Both compounds are competitive inhibitors that bind to the ATP binding pocket of these kinases to prevent their catalytic activity [30,31]. Both inhibitors were prepared in DMSO, and added 1 h before CPT treatment. The proteasome inhibitor MG132 (Bio-Techne) was also prepared in DMSO and applied at 10 µM 1 h prior to CPT treatment. In all cases, control cultures were treated with an equivalent volume of DMSO. Total DMSO concentration was always below 0.5 %.

### RNA extraction and cDNA synthesis

N2a neurons were washed with phosphate buffered saline (PBS) and total RNA was then extracted with Qiazol (Qiagen) following the instructions of the manufacturer. The concentration of RNA in each sample was then measured using NanoDrop One UV-Vis (ThermoFisher Scientific), and 5 µg of total RNA were used for the synthesis of the cDNA in a 20 µl reaction mix containing 4 µl of 5x reaction buffer (Invitrogen), 40 units of RNAsin (Promega), 250 ng/µl of random primers (Invitrogen), 10 mM dNTPs (Biotools), 10 mM dithiothreitol (DTT) (Invitrogen), and 200 units of SuperScript IV (Invitrogen), following the instructions of the reverse transcriptase manufacturer.

### Quantitative RT-PCR (qRT-PCR)

qRT-PCR was performed with the QuantStudio 3 Real-Time PCR system (Thermo Fisher Scientific) using SYBR Green (Cultek) in 15 µl reaction volume containing 1 µl cDNA and the upstream and downstream oligonucleotides (see Supplementary Table 3) at 10 µM.

### Cell extracts

Cultures were placed on ice, washed with ice-cold PBS, and incubated for 3 min with 200 μl of cold extraction buffer containing 20 mM Tris-HCl (Roche) pH 6.8, 1 mM EDTA (Merck), 1% Triton-X100 (Sigma), 0.5% sodium dodecyl sulfate (Sigma-Aldrich), 10 mM β-mercaptoethanol (Sigma-Aldrich), and 1× cOmplete Mini, and EDTA-free, protease inhibitor cocktail (Roche). 1× Phosphatase Inhibitor Cocktail 2 (Sigma-Aldrich) was added to the cell lysing buffer in experiments where preservation of phosphoproteins was mandatory. Cell extracts were then scraped with a rubber policeman and homogenized using a 22G needle. Protein was then quantified using the Protein Assay Dye Reagent Concentrate system (BioRad).

### Subcellular fractionation

Subcellular fractionation was performed using a modification of the protocol described by [32]. Briefly, cultures were placed on ice, washed with ice-cold PBS, and lysed with 500 μl of cold lysis buffer containing 10 mM HEPES (Sigma-Aldrich) pH 7.6, 10 mM NaCl (Merck), 1 mM KH_2_PO_4_ (Merck), 5 mM NaHCO_3_ (Merck), 5 mM EDTA (Merck), 1× cOmplete Mini, and EDTA-free, protease inhibitor cocktail. 1× Phosphatase Inhibitor Cocktail 2 was added to the cell lysing buffer in experiments where preservation of phosphoproteins was mandatory. Cell lysates were then scraped with a rubber policeman and homogenized by 10 passages through a 22G needle, and an aliquot of the cell lysate was kept as a lysate control. Cell nuclei were spun down at 300 × g for 10 min and the nuclear pellet was kept at 4 °C. The supernatant containing membranes and cytosol was then microfuged at 16,000 × g for 45 min at 4°C. The supernatant (cytosol fraction) was removed and concentrated with Vivaspin 500 Centrifugal Concentrator filters (cutoff, 3 kDa) (Sartorius) as described by the manufacturer. In addition, lysis buffer containing 1% Triton X-100 was added to the pellet (membrane fraction). Finally, nuclear proteins were extracted for 15 min under frequent vortexing in high-salt buffer containing 5 mM HEPES pH 8.0, 420 mM NaCl, 0.25 mM MgCl_2_ (Merck), 0.1 mM EDTA, 12.5% glycerol (PanReac), 1× cOmplete Mini, EDTA-free, protease inhibitor cocktail, and 1× Phosphatase Inhibitor Cocktail 2. The extract was then centrifuged at 16,000 × g for 10 min at 4°C and the supernatant (soluble nuclear fraction) was kept on ice.

### p38^MAPK^ and JNK assays

A luminiscent p38^MAPK^ assay was performed using the p38α Kinase Enzyme System (Promega) and the ADP-Glo kinase Assay kit (Promega). The JNK1 Kinase Enzyme System (Promega) together with the ADP-Glo kinase Assay kit was used to evaluate the activity of JNK. TAT and TAT-E2F4DN peptides were dissolved at 10 mM in 1× kinase reaction buffer [40 mM Tris-HCl pH 7.5, 20 mM MgCl_2_, 50 micro-M DTT and 0.4 μg/μl bovine serum albumin (Sigma-Aldrich)]. Phosphorylation reactions (5 μl final volume) were performed in Nunc 384-well white plates (ThermoFisher Scientific) containing 10 ng of p38α (or of JNK1), 200 ng of the p38α (or of JNK1) substrate provided by the manufacturer, and the TAT or TAT-E2F4DN peptides at the indicated concentrations. Kinase reactions were initiated by adding ATP at a final concentration of 30 μM. After 30 min at room temperature (RT), 5 μl of ADP-Glo reagent was added and incubated at RT for 40 min to stop the kinase reactions. Then 10 μl of Kinase Detection Reagent was added and incubated for 30 min at RT. The light generated was measured with a BMG FLUOstar OPTIMA Microplate reader (BMG Labtech).

### Dephosphorylation assay

A cell extract volume containing 16 µg of protein was incubated for 2 h at 37 °C in 200 µl 1× FastAP reaction buffer (ThermoFisher Scientific) containing 100 U FastAP thermosensitive alkaline phosphatase (AP) (ThermoFisher Scientific). Then, the reaction was stopped by adding 50 µl of 250 mM EDTA. Finally, reactions were concentrated with Vivaspin 500 Centrifugal Concentrator filters (cutoff, 3 kDa), as described by the manufacturer.

### Western blotting and image analysis

Cell extracts and subcellular fractions obtained as described above or AP-treated proteins were boiled for 5 min in 1× Laemli buffer. The extracts were fractionated by SDS PAGE on acrylamide gels of the appropriate concentration, and transferred to 0.45 µm Immobilon-FL polyvinylidene difluoride (PVDF) transfer membranes (Millipore). The membranes were incubated for 1 h with Intercept (TBS) Blocking Buffer solution (LI-COR) (IBB), and then incubated overnight at 4 °C with the appropriate antibody in IBB. After washing the membranes three times in TBS containing 0.1% Tween 20 (TBS-T), they were incubated for 1 h at RT with a 1/15,000 dilution of secondary antibodies in IBB containing 0.1% Tween 20. Finally, they were washed again with TBS-T as described above, followed by a final wash with TBS. The protein bands were then visualized using the Odyssey CLx Infrared Imaging System (LI-COR). Gels were overrun to resolve the E2F4 phosphorylation bands (60-70 kDa), thus resulting in the loss of the Histone H4 band due to its extreme high mobility (15 kDa). Therefore, different gels were used to obtain the E2F4 bands and histone localization controls. Band intensities were quantified using LicorLite using the “asign shape” command for background subtraction. The intensity for each E2F4-specific band obtained in subcellular fractions was quantified using ImageJ software. The intensity of bands corresponding to proteins of interest were normalized to the intensity of actin or lamin B1 in the same lane.

### Immunocytochemistry

Immunocytochemistry was performed in N2a neurons cultured on glass coverslips fixed for 15 min at RT with 4% paraformaldehyde (PFA; Merck). After three washes with PBS, cells were permeabilized and blocked with PBS containing 0.1% Triton-X100 (PBS-T) supplemented with 10% fetal bovine serum (FBS) (blocking medium) for 1 h. Coverslips were then incubated for 2 h at RT with blocking medium and the appropriate primary antibody. After three washes in PBS-T, coverslips were incubated for 1 h in blocking medium containing the appropriate secondary antibody and 0.1 μg 4’,6-diamidine-2’-phenylindole (DAPI; Sigma-Aldrich). After three washes with PBS-T plus one additional washing step with PBS, coverslips were mounted with InmunoSelect Antifade Mounting Medium (Dianova) on glass slides. Images of the cells were taken by confocal microscopy.

### Confocal microscopy and image analysis

Confocal images were acquired with a Leica SP5 inverted confocal microscope. In all analyses, a range of 10-35 images of 1 µm thick were taken per stack at 40× magnification so that the entire thickness of the cells present in the microscope field is included, and then the maximum projection was obtained. Images used for the analysis of γH2AX were taken at 40× magnification with 2× zoom.

Image analysis was performed using ImageJ (Fiji). For the analysis of caspase-3 activation, the number of cells positive for caspase-3 cleaved at Asp175 were counted and quantified as the percentage of DAPI labeling. For the analysis of DNA damage foci through γH2AX immunostaining, cells were classified based on the nuclear labeling pattern as well as the number of foci present in their nuclei. For the analysis of the phosphoE2F4 vs. E2F4 ratio, DAPI was used as a reference for cell identification and the same setting was used for both phosphoE2F4 and E2F4 labeling channels. Briefly, after adjusting the cell outlines using the “threshold” command, the “fill holes” and the “watershed” commands were used to improve the selection of the cells. Then, the region of interest (ROI) tool was used to measure the “integrated density” of each cell, which was subtracted from the average value of the background signal outside the cells. ImageJ software was used for the analysis of the 6xHis immunofluorescence levels using the rectangle tool. The pixel intensity mean was quantified for each non-transfected cell to set up the background threshold. Cells were considered to be positive for the staining if they showed mean values above this threshold.

### Statistical analysis

Statistical analyses were carried out using GraphPad Prism software (version 9). Quantitative data are represented as the mean ± standard error of the mean (SEM). Statistical tests used in this study and the number of independent samples for each experimental group are specified in the figure legends. Differences were considered to be statistically significant with a p-value < 0.05.

## Results

### Low serum concentration favors neuronal differentiation of N2a neuroblastoma cells

To study the role of E2F4 phosphorylation in neuronal DNA damage response we focused on N2a neuroblastoma cells due to the capacity of this cell line to differentiate as neurons under low serum conditions [33–36]. N2a cells were therefore cultured in the presence of 1% FBS to induce neuronal differentiation [36]. Under these conditions, N2a cells increased the number of processes resembling dendrites (Supplementary Fig. 1A), which were immunopositive for Microtubule-associated Protein 2 (MAP2) (Supplementary Fig. 1B), known to be located in the dendrites and cell bodies but not axons from neurons [37]. Moreover, these cells (which will be hereafter referred to as N2a neurons) expressed both *Syn1*-specific mRNA (Supplementary Fig. 1C) and Synapsin1 protein (Supplementary Fig. 1B). Nevertheless, in contrast to the study by [38], which showed increased *Syn1* expression in differentiated N2a neuroblastoma, we observed that this protein and its coding mRNA was expressed at similar levels in both undifferentiated N2a neuroblastoma cells and N2a neurons (Supplementary Fig. 1B,C). In any case, the capacity to express Synapsin 1 makes these cells an ideal system to express exogenous proteins under the control of the human synapsin1 (*hSyn1*) promoter [39].

### Expression of E2F4 and its mutant forms E2F4DN and E2F4CA in N2a neurons

We confirmed that N2a neurons were able to express E2F4WT and its mutant forms E2F4DN and E2F4CA, driven by the *hSyn1* promoter. By using an anti-6xHis specific antibody, we verified by immunocytochemistry that E2F4WT-6xHis, E2F4DN-6xHis and E2F4CA-6xHis are widely expressed in N2a neurons 48 hours after transduction with the Ad5.E2F4WT-6xHis, Ad5.E2F4DN-6xHis, and Ad5.E2F4CA-6xHis vectors, respectively (Fig. 1A). Non-transduced control cells showed unspecific labeling that allowed us to set up a threshold above which a cell was classified as positive (Supplementary Fig. 2). Quantification of the percentage of 6xHis-positive cells based on this threshold indicated that above 95% of N2a neurons expressed the aforementioned constructs (Fig. 1B). Moreover, western blot (WB) analysis with the anti-6xHis antibody confirmed the expression of these proteins at physiological levels, as they show intensities comparable to that of actin (Fig. 1C). Interestingly, the different forms of E2F4 displayed a multiband pattern (Fig. 1C). This suggests that, in N2a neurons, E2F4 can be phosphorylated in multiple residues, as previously described [17,40]. As in these later studies, we found five different bands for this transcription factor (bands *a* to *e*), with different mobility. As expected, the phosphomimetic nature of E2F4CA led to the presence of lower mobility bands, which were also named from *a* to *e*, being bands *b* and c those with the highest intensity (Fig. 1C).

**Fig. 1.**
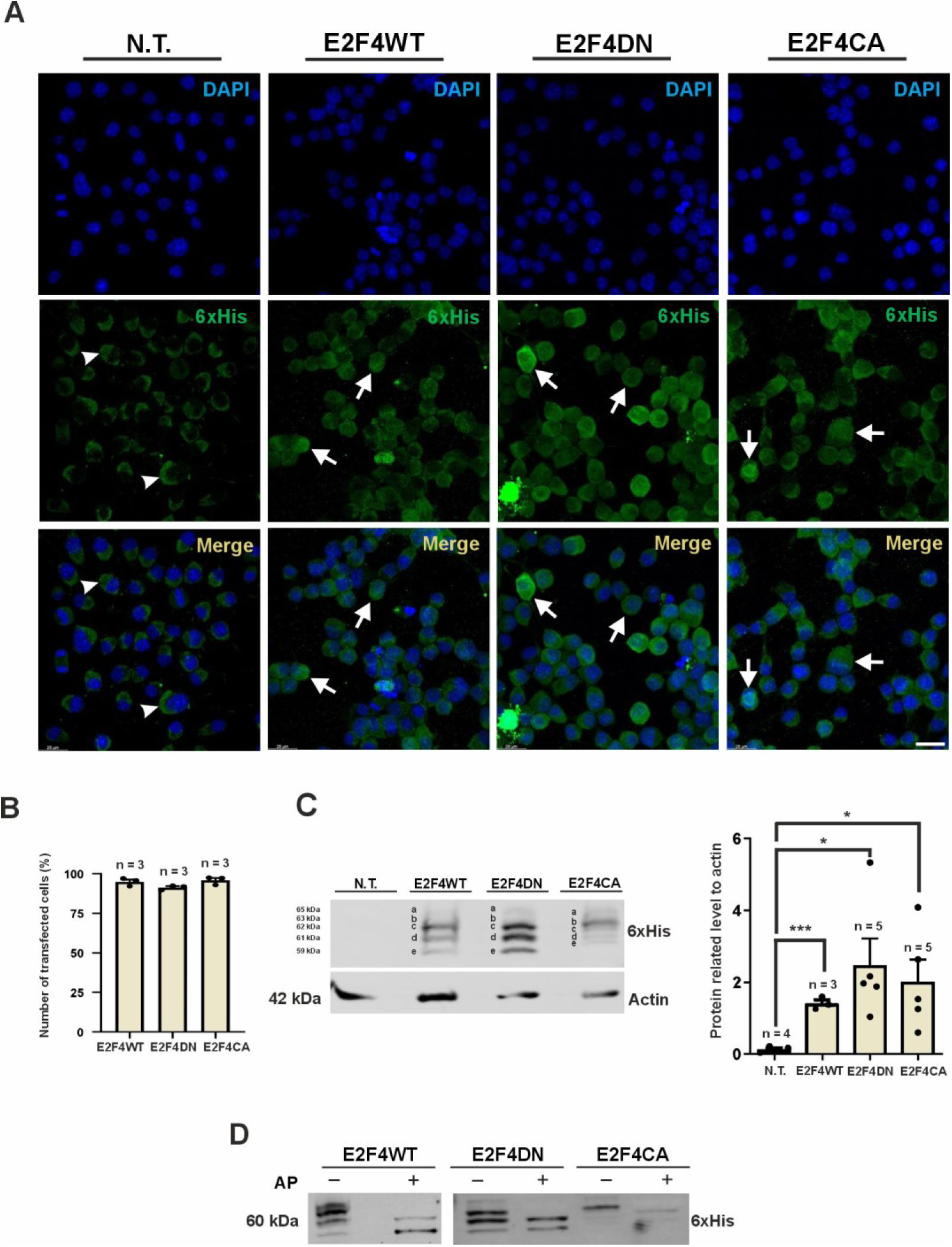
Overexpression and phosphorylation of different E2F4 forms in N2a neurons. **A** Representative inmunostaining of N2a neurons, either non-transfected (N.T.) or transduced with Ad5 vectors expressing E2F4WT-6xHis (E2F4WT), E2F4DN-6xHis (E2F4DN) or E2F4CA-6xHis (E2F4CA), using an anti-6xHis antibody (green). Notice that, while unspecific fluorescence can be observed in the non-transfected cells (arrowhead), characterized by a crescent-like structure that is not observed in transduced cells, 6xHis-specific labeling with different intensity levels was observed in the transfected cells (arrows). Cell nuclei were labeled with DAPI (blue). Bar: 25 μm. **B** Quantification of E2F4WT-6xHis, E2F4DN-6xHis and E2F4CA-6xHis transduction efficiency in the N2 neuronal cultures shown in a (i.e. number of 6xHis+ cells related to DAPI+ nuclei). **C** Left panel: representative western blot of the multiband pattern (bands a-e) observed in cell extracts from E2F4WT-6xHis, E2F4DN-6xHis or E2F4CA-6xHis-transduced N2a neurons, as revealed with an anti-6xHis antibody. Actin was used as a loading marker. Right panel: quantification of the sum of intensities for the different bands shown in the left panel relativized to actin. *p<0.05; ***p< 0.001 (unpaired Student’s *t* test). **D** Representative western blots of the effect of alkaline phosphatase (AP) on the multiband pattern observed in cell extracts from E2F4WT-6xHis-, E2F4DN-6xHis- or E2F4CA-6xHis-transduced N2a neurons.

The incubation of the cell extracts with AP demonstrated that most of the bands that were observed in E2F4WT-, E2F4DN-, and E2F4CA-expressing N2a neurons derive from different phosphorylation states of the molecule. In this regard, only two high mobility bands (*d* and *e*) remained in the presence of this enzymatic activity (Fig. 1D), suggesting that one of these two bands contains a post-translational modification different from phosphorylation [8]. Interestingly, the intensity of band *e* increased upon AP treatment in extracts from E2F4WT-expressing cells, suggesting that this latter band contains the modified form of E2F4 in unphosphorylated state, which reduces its mobility to become part of band *d* upon phosphorylation. This hypothetical modified form of E2F4 seems to become phosphorylated in the Thr248/Thr250 motif since the incubation of the E2F4DN-derived extracts with AP does not increase the intensity of band *e* in comparison to band *d* (Fig. 1D), while the incubation of the E2F4CA-derived extracts with this enzyme leads to an enrichment of band *d* (Fig. 1D).

To gain further insight into the nature of band *e*, E2F4DN-, E2F4CA-, and E2F4WT-transduced N2a neurons were treated with the proteasome inhibitor MG132 and then either kept for 9 h or treated with CPT 1 h later and maintained for 8 additional hours before obtaining cell extracts. The inhibition of the proteasome resulted in the loss of band *e* and the enrichment of band *b* in N2a neurons expressing E2F4WT or E2F4DN in the absence of CPT (Supplementary Fig. 3). These results suggest that the degradation by the proteasome of an upstream regulator that indirectly targets the phosphorylated form of E2F4 migrating as band *b* is crucial for the generation of band e.

Our analysis also indicated that E2F1 expression in N2a neurons results in a multiband pattern (Supplementary Fig. 4), which is consistent with the phosphorylation of this protein in several residues [41,42].

### Dynamic localization of E2F4WT, E2F4DN and E2F4CA in subcellular domains

To analyze the cell compartments where exogenous E2F4WT, E2F4DN, and E2F4CA locate in N2a neurons, we obtained subcellular fractions including membrane, cytosol and nucleus. This analysis indicated that, as expected from a soluble protein, all forms of E2F4 were present in the cytosolic and nuclear compartments while E2F4 was absent from the membrane fraction (Fig. 2A). The purity of the cytosolic fraction was confirmed by the presence of a single band of GAPDH of ∼36 kDa (Supplementary Fig. 5). In addition, the nuclear fraction specifically contained Histone H4 and two GAPDH-specific bands, a faint band showing equivalent mobility as in the cytosol (∼36 kDa) and another prominent band of higher mobility (∼34 kDa) [43], while the membrane fraction specifically contained the sodium channel (Supplementary Fig. 5), thus confirming the purity of these subcellular fractions. The pattern of band intensities differed between the nuclear fraction and the cytosolic fraction. Bands *c* and *d* were the main phosphorylated states observed in the cytosol for both E2F4WT and E2F4DN, while E2F4CA, in accordance with its phosphomimetic nature, displayed higher levels of phosphorylation in this subcellular compartment as its main bands were *b* and *c* (Fig. 2B). In contrast, a shift towards bands with increased phosphorylated levels was detected in the nucleus, with different degrees depending on the specific form of E2F4 that was analyzed. Nuclear E2F4CA was mainly in the form of band *b*, while the predominant state of E2F4WT was band *c*. Finally, bands *b*, *c*, and *d* were equally displayed for nuclear E2F4DN, indicative of its reduced capacity to become phosphorylated due to the Thr248Ala/Thr250Ala mutation. The differences of phosphorylation state of E2F4DN and E2F4CA (i.e. higher phosphorylation in E2F4CA vs lower phosphorylation in E2F4DN) suggest that the function of E2F4 as a transcription factor, but also as a homeostatic protein [8], may differ depending on the phosphorylation state of this latter Thr motif.

**Fig. 2.**
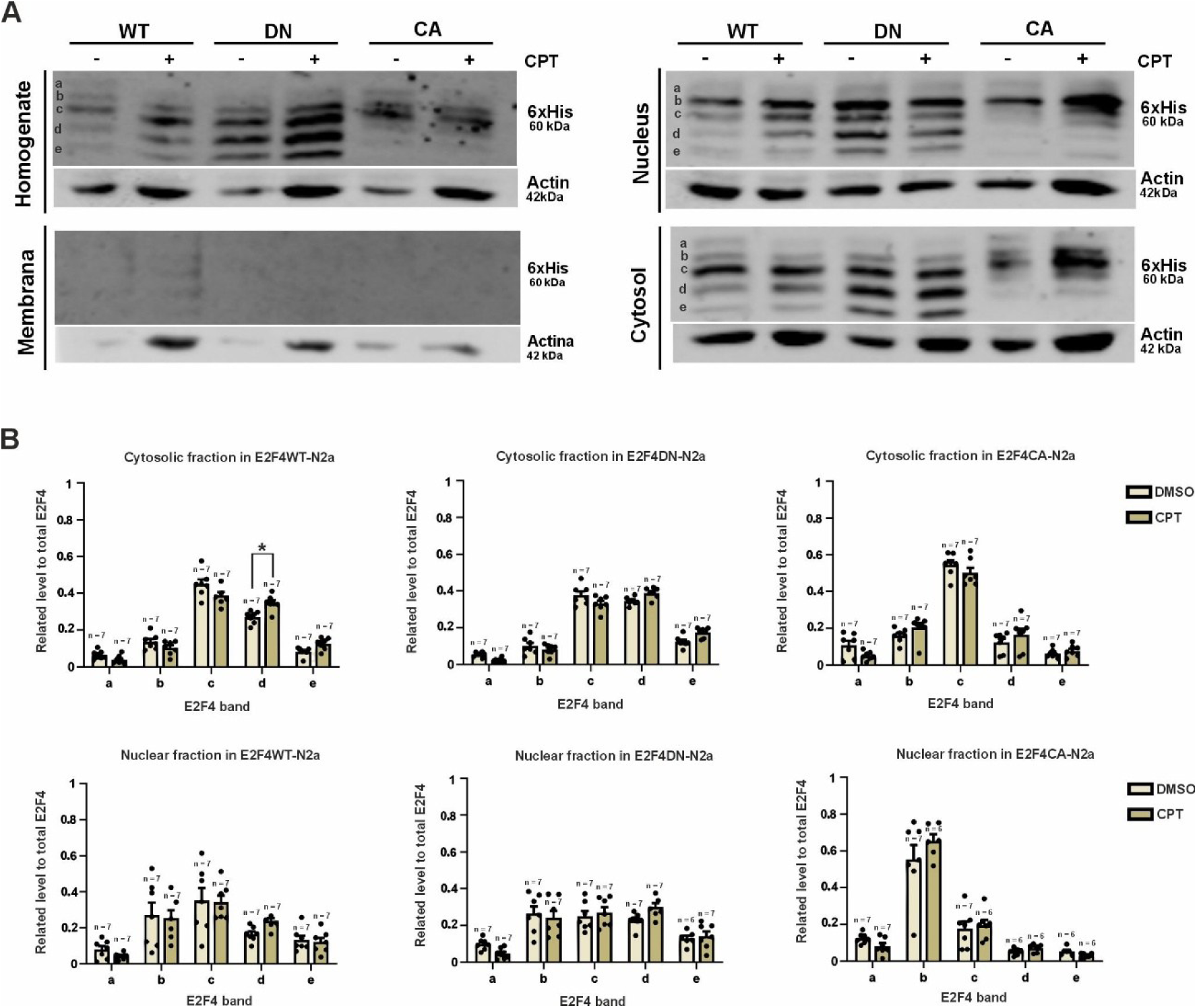
Subcellular localization of the different phosphorylation forms of E2F4 in N2a neurons. **A** Representative western blots of the multiband pattern detected in the indicated homogenates and subcellular fractions from E2F4WT-6xHis- (E2F4WT), E2F4DN-6xHis- (E2F4DN) or E2F4CA-6xHis-transduced N2a neurons (E2F4CA), either under control conditions or 8 h after CPT treatment, as revealed with an anti-6xHis antibody. Actin was used as a loading marker. **B** Quantification of the relative frequency of the bands a-e that are present in the cytosolic and nuclear fractions for each experimental condition (see panel A). *p<0.05 [Two-way ANOVA, followed by Tukey’s honestly significant difference (HSD) post-hoc test].

To study whether DNA damage may induce specific E2F4 phosphorylation at the Thr248/Thr250 motif, we focused on N2a neurons treated with 10 µM CPT as a paradigm of DNA damage [28]. The presence in the culture medium of this topoisomerase I inhibitor for 8 h did not substantially modified the phosphorylation profile observed in N2a neurons (Fig. 2A, B), with the exception of band *d* in the cytosolic fraction from E2F4WT-expressing N2a neurons, which significantly increased its intensity in the presence of CPT (Fig. 2B). The intensity of this band was not significantly altered in the cytosolic fraction of both E2F4DN- and E2F4CA-expressing N2a neurons, suggesting that it results from the phosphorylation of the Thr248/Thr250 motif of E2F4 in response to CPT treatment.

### CPT treatment induces p38^MAPK^ activity in N2a neurons

In chick neurons, the stress kinase p38^MAPK^ can phosphorylate E2F4 in the Thr265/Thr267 conserved domain [16], which is orthologous of Thr248/Thr250 in human E2F4 and Thr249/Thr251 in mouse E2F4 (notice that we will refer to Thr248/Thr250 in those experiments where the different variants of human E2F4 are expressed using Ad5 vectors, whereas the term Thr249/Thr251 will be used for those experiments where the phosphorylation state of endogenous mouse E2F4 is evaluated). In a mouse model of AD, E2F4 has been shown to become phosphorylated in Thr249 [9], likely being this phosphorylation part of the etiopathology of the disease, which includes p38^MAPK^ activation at its early stages [44–47]. These observations suggest that stressful conditions triggering p38^MAPK^ activation may lead to p38^MAPK^-dependent phosphorylation of E2F4. To directly test the hypothesis that E2F4 can be phosphorylated by p38^MAPK^ in the Thr249/Thr251 motif, N2a neurons were treated with 10 µM CPT to induce DNA damage, as a mechanism of cell stress followed by cell death in neurons [23]. We found that, under these conditions, p38^MAPK^ activity becomes gradually activated, an effect that is already evident at 6 h while showing statistically significant differences 8 h after CPT treatment (Fig. 3A, B). This effect, which was based on the dual phosphorylation of p38^MAPK^ on Thr180 and Tyr182 [48], was specific for p38^MAPK^ since the levels of active c-Jun N-terminal kinases (JNK) were not significantly upregulated at the studied time points (Fig. 3C, D).

**Fig. 3.**
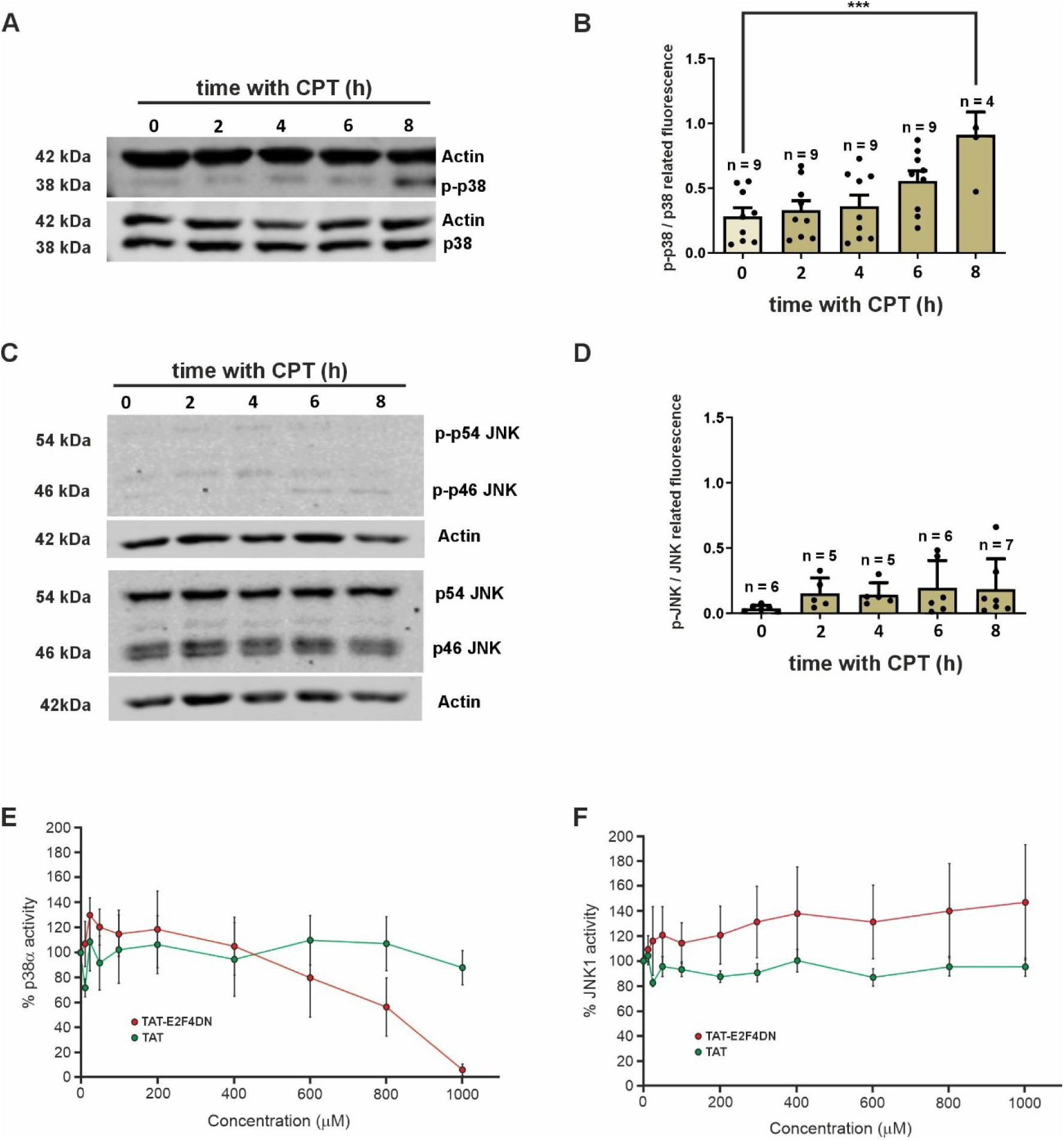
p38^MAPK^ activation in response to CPT in N2a neurons. **A** A representative western blot from cell extracts of N2a neurons isolated at the indicated time points after 10 µM CPT treatment, using an antibody recognizing active p38^MAPK^ (p-p38) and total p38^MAPK^ (p38). Actin was used as a loading marker. **B** Quantification of p38^MAPK^ activation as the ratio between active p38^MAPK^ (relativized to Actin) and total p38^MAPK^ (relativized to Actin) at the indicated time points after CPT treatment. **p<0.01; ***p<0.001 (One-way ANOVA, followed by Tukey’s HSD post-hoc test). **C** Representative western blot from cell extracts of N2a neurons isolated at the indicated time points after 10 µM CPT treatment, using an antibody recognizing active JNK (p-p54 JNK and p-p46 JNK) and total JNK (p54 JNK and p46 JNK). Actin was used as a loading marker. **D** Quantification of JNK activation as the ratio between the sum of active JNK (p-p54 JNK plus p-p46 JNK relativized to Actin) and total JNK (p54 JNK plus p46 JNK relativized to Actin) at the indicated time points after CPT treatment. **E** A p38^MAPK^ kinase assay performed in the presence of either a control peptide (TAT) or a TAT-E2F4DN peptide at the indicated concentrations. **F** A JNK assay performed in the presence of either a control peptide (TAT) or a TAT-E2F4DN peptide at the indicated concentrations.

### p38*^MAPK^* specifically phosphorylates Thr249 in endogenous E2F4

To verify that the activation of p38^MAPK^ in N2a neurons treated with 10 µM CPT results in the phosphorylation of endogenous E2F4 in its Thr249/Thr251 motif, we used an antibody that recognizes the phosphoThr249 epitope of mouse E2F4, previously generated in our laboratory [9]. To this aim, N2a neurons were cultured in the presence of 10 µM CPT for 0 h (untreated control cells), 4 h, 6 h and 8 h. Under these conditions, E2F4 expression levels were increased (Supplementary Fig. 6). These cultures were then fixed and immunostained with the phospho-specific anti-E2F4 antibody together with an antibody recognizing E2F4 independently of its phosphorylation state, and then the phosphoE2F4/E2F4 ratio was calculated for each experimental condition. We found that the phospho-E2F4/E2F4 ratio was significantly higher in N2a neurons treated with CPT for 8 h (Fig. 4A, B), in accordance with the significant activation of p38^MAPK^ at this time point (see above). This result is consistent with the significant increase of the band *d*, observed specifically in the cytosol of E2F4WT-, but not E2F4DN- or E2F4CA-expressing N2a neurons treated for 8 h with CTP (Fig. 2A, B).

**Fig. 4.**
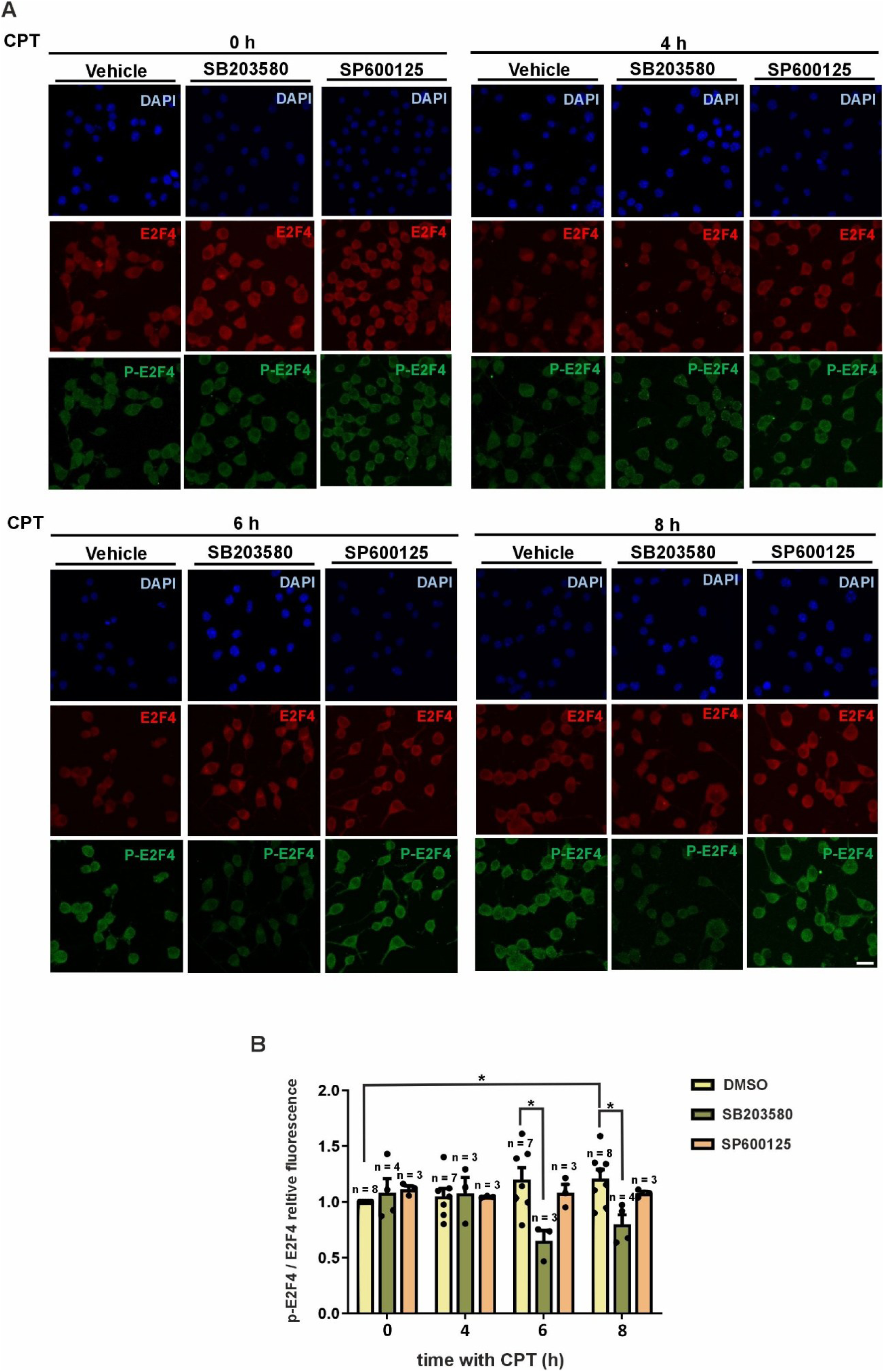
E2F4 phosphorylation at Thr249 in N2a neurons treated with CPT. **A** Representative immunostaining of N2a neurons treated with 10 µM CPT for the indicated time points in the presence of vehicle, the p38^MAP^ inhibitor SB203580 or the JNK inhibitor SP600125, using antibodies specific for total E2F4 (E2F4) or E2F4 phophorylated at Thr249 (P-E2F4). Cell nuclei were labeled with DAPI (blue). Bar: 25 μm. **B** Quantification of endogenous E2F4 phosphorylation at Thr249 of in response to 10 µM CPT treatment by measuring the ratio between p-E2F4 and E2F4 fluorescence intensities for each experimental condition. *p<0.05 (Two-way ANOVA, followed by Tukey’s HSD post-hoc test).

To test whether the Thr249-specific phosphorylation of E2F4 in response to CPT was associated to p38^MAPK^ activity, N2a neurons were cultured in the same conditions as above either in the presence of 10 µg/µl SB203580, a p38^MAPK^-specific inhibitor [49,50], or in the presence of 25 µg/µl SP600125, a JNK-specific inhibitor [31], the latter as a control. This analysis indicated that the presence of SB203580, but not SP600125, was able to significantly reduce the phosphorylation of E2F4 at both 6 h and 8 h (Fig. 4A, B), the time points at which p38^MAPK^ activation in response to the CTP treatment was already evident (Fig. 3A, B).

A cell-free kinase assay was performed to confirm the capacity of p38^MAPK^ to functionally interact with E2F4. By incubating a p38^MAPK^-specific substrate together with ATP and activated p38α, we were able to show dose-dependent inhibition of the p38^MAPK^ activity using a mouse E2F4-derived peptide containing the Thr249Ala/Thr251Ala mutation (Fig. 3E). In contrast, this same peptide was not able to prevent the JNK activity when tested in a cell-free JNK1 kinase assay (Fig. 3F).

### E2F4DN expression prevents CPT-induced cell death in N2a neurons

CPT treatment is known to induce apoptosis [23], which is potentiated by E2F1 expression in a cell cycle-independent manner [2,24,25]. Therefore, we performed a temporal analysis to evaluate the time point at which N2a neurons overexpressing E2F1 start to die in response to CPT, as evidence by caspase-3 activation. This analysis indicated that a statistically significant increase of apoptosis can be detected in these cells 48 h after CPT treatment (Supplementary Fig. 7).

CPT treatment can lead to the degradation of Rb protein and phosphorylation of its family members, p107 and p130 [51,52]. Therefore, the release of E2F4 may participate in CPT-induced apoptosis, as all these pocket protein members are known E2F4 interactors [53]. To test whether E2F4 and its phosphorylation at the Thr248/Thr250 motif participates in CPT-induced apoptosis, we cultured N2a neurons transduced with E2F4CA, followed by CPT treatment for 24 h or 48 h. This analysis indicated that, like E2F1, E2F4CA was able to induce a statistically significant increase of apoptosis in N2a neurons when co-treated with 10 μM CPT for 48 h, but not 24 h (Fig. 5A,B). In contrast, no significant induction of apoptosis was observed in these cells in the presence of E2F4DN (Fig. 5A,B). Therefore, the phosphorylation of E2F4 in its Thr248/Thr250 motif is likely required for the trigger of apoptosis by CPT.

**Fig. 5.**
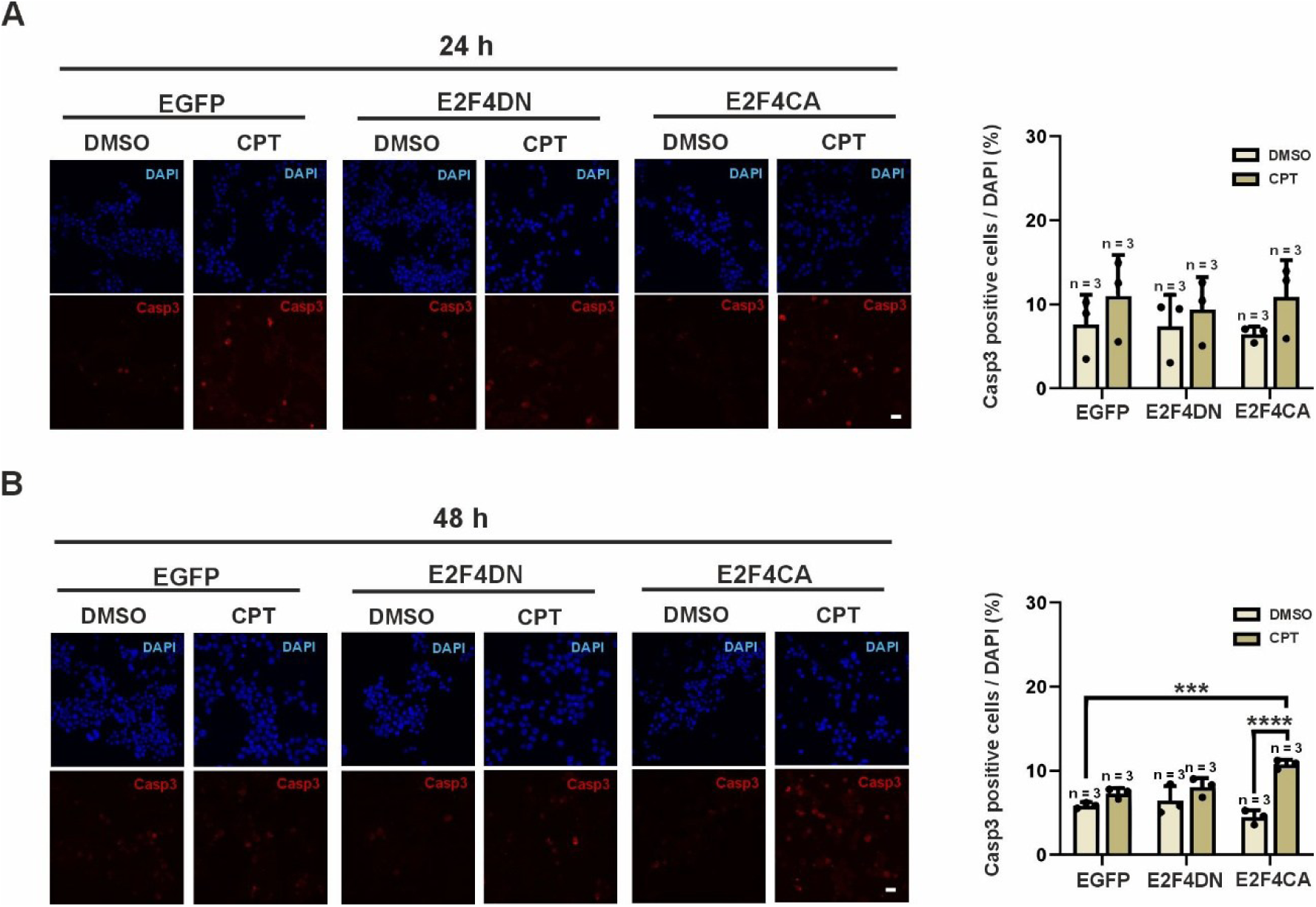
Regulation of apoptosis by E2F4 phosphorylation in CPT-treated N2a neurons. Percentage of active caspase-3-positive N2a neurons related to total DAPI-positive nuclei for each indicated experimental condition. ***p<0.005; ****p<0.0001 (Two-way ANOVA, followed by Tukey’s HSD post-hoc test).

### The antiapoptotic effect of E2F4DN in N2a neurons is independent of DNA repair

DNA damage is known to trigger apoptosis, mainly through a p53-dependent mechanism [54] although it may also rely on the transfer RNAse SLFN11 in p53 mutant cells [55]. Therefore, the antiapoptotic effect of E2F4DN in CPT-treated N2a neurons might depend primarily on the potential effect of E2F4 to favor DNA repair since it regulates the expression of DNA repair genes [8]. Alternatively, E2F4 might prevent apoptosis in these cells by directly acting on the proapoptotic machinery downstream of DNA damage. To test whether E2F4DN prevents DNA damage in response to CPT treatment prior to the detection of apoptosis, N2a neurons, previously transduced with EGFP, E2F4DN or E2F4CA, or left untreated, where maintained for either 0, 8, or 32 h in the presence of vehicle or of 10 μM CPT. Then, cultures were fixed and immunostained with an anti-γH2AX antibody to identify double strand DNA breaks [56]. This analysis demonstrated that DNA damage is increased with time for all the experimental conditions that were analyzed, as revealed by the γH2AX-specific labeling. Indeed, while the proportion of cells with punctate γH2AX labeling (i.e. cells with low DNA damage levels) gradually decreased with time (Fig. 6A, C), those showing a diffuse γH2AX pattern (i.e. cells with strong DNA damage) progressively increased its proportion with time (Fig. 6A, D).

**Fig. 6.**
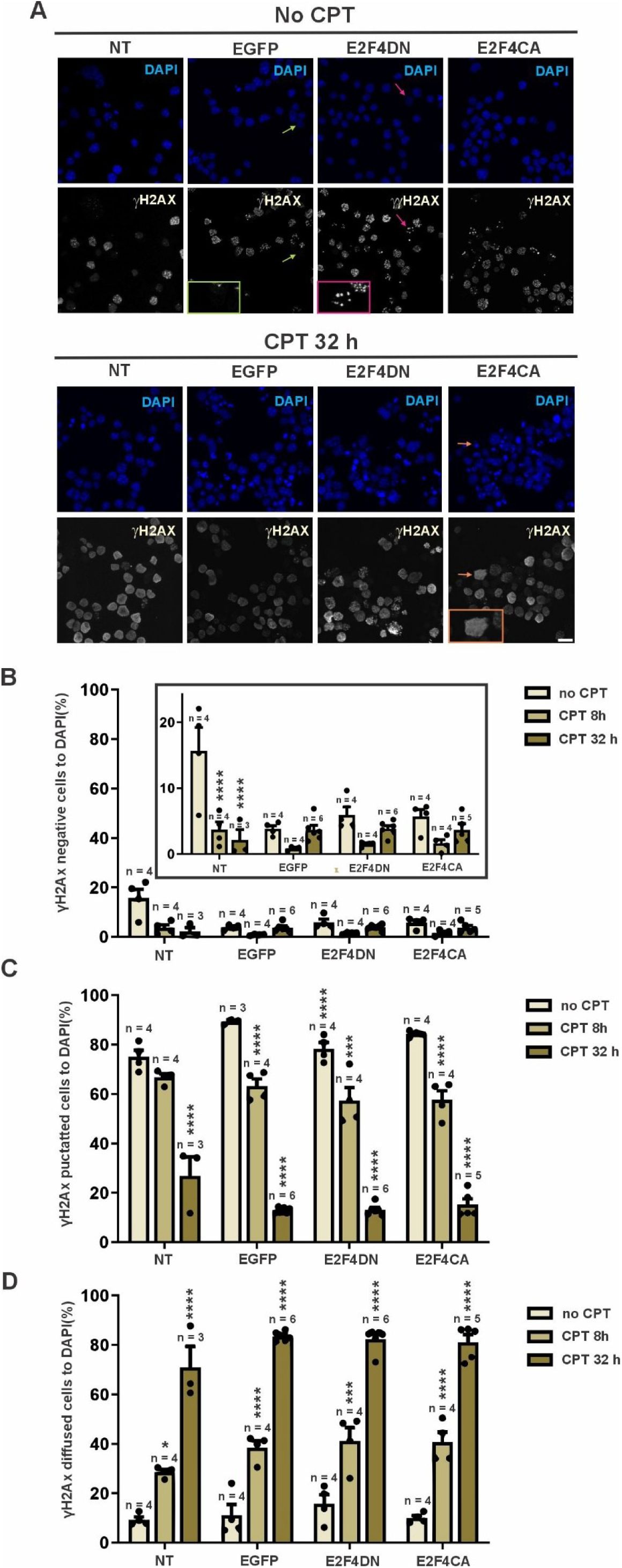
DNA damage regulation by E2F4 in N2a neurons. **A** Representative inmunostaining performed with an antibody specific for γH2AX to identify double strand breaks in the genomic DNA of N2a neurons transduced with either EGFP (EGFP), E2F4DN-6xHis (E2F4DN) or E2F4CA-6xHis (E2F4CA) and treated with 10 µM CPT for 32 h or left untreated. Orange arrow: a γH2AX-positive cell with diffuse staining shown at high magnification in the orange box. Green arrow: a γH2AX-negative cell shown at high magnification in the green box. Pink arrow: a γH2AX-positive cell with punctate staining shown at high magnification in the pink box. Bar: 25 μm. **B** Quantification of the γH2AX-negative N2a neurons, related to DAPI-positive nuclei, for each experimental group. **C** Quantification of the γH2AX-positive N2a neurons showing punctate staining, related to DAPI-positive nuclei, for each experimental group. **D** Quantification of the γH2AX-positive N2a neurons showing diffuse staining, related to DAPI-positive nuclei, for each experimental group. *p<0.05, ***p<0.005, ****p<0.001 (Two-way ANOVA, followed by Tukey’s HSD post-hoc test).

Interestingly, no statistically significant differences were observed at any time point in the proportion of CPT-treated N2a neurons either lacking γH2AX-specific immunostaining or showing punctate or diffuse γH2AX labeling when E2F4DN or E2F4CA were expressed as compared to control cells (Fig. 6B-D). Therefore, our results indicate that the antiapoptotic effect of E2F4DN in response to CPT-induced DNA damage likely relies on its interaction with the downstream pathways involved in cell death.

### Cited2 acts downstream of E2F4DN to prevent CPT-dependent N2a neuronal death

Cited2 is a transcriptional regulator that can be regulated by the crosstalk of E2F1 and E2F4 in response to CPT treatment [26]. Depending on the cellular context, this protein can act either as an antiapoptotic factor [57,58], since it can prevent p53 activation [59], or as a proapoptotic agent [60,61]. Therefore, we hypothesized that Cited2 participates in the control of CPT-dependent cell death by E2F4DN due to the capacity of the latter to induce Cited2 expression. This hypothesis was tested using E2F4DN- and E2F4CA-expressing N2a neurons treated for 4 h with CPT to avoid any interference with active p38^MAPK^ since, at this time point, p38^MAPK^ has not yet been activated. This analysis demonstrated that E2F4DN upregulates *Cited2* gene expression 4 h after CPT treatment when compared to both 0 h and 2 h. This contrasts with the inability of E2F4CA to significantly induce *Cited2* expression at this time point (Fig. 7A), thus indicating that Thr248/Thr250 phosphorylation prevents the capacity of E2F4 to express *Cited2*. Under control conditions, *Cited2* expressions was slightly upregulated 2 h after CPT treatment, but its levels of expression did not increase at the 4h time point.

**Fig. 7.**
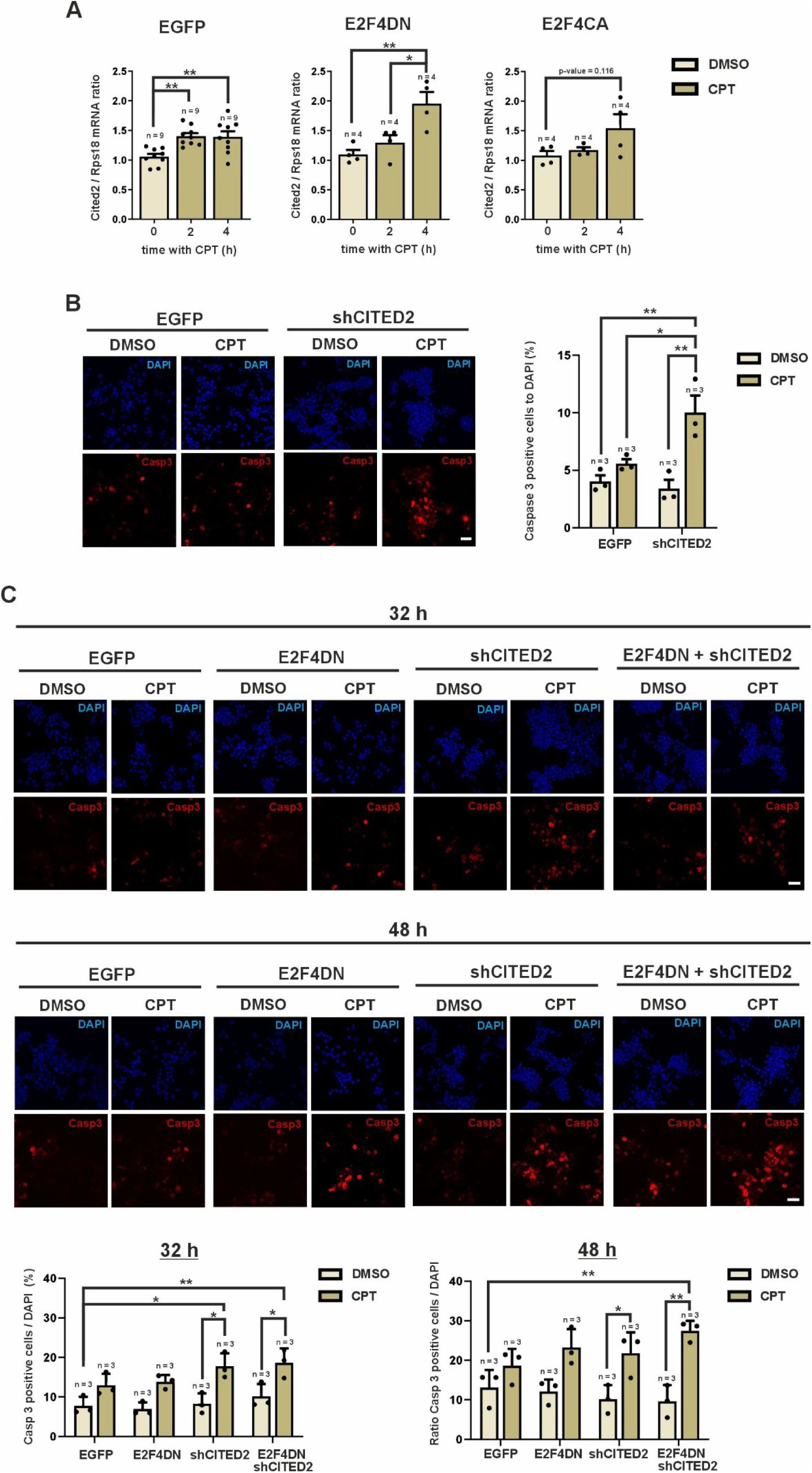
Participation of Cited2 in the CPT-induced apoptotic cell death regulated by E2F4 in N2a neurons. **A** Quantification of *Cited2* mRNA levels by real-time quantitative PCR (RT-qPCR), normalized to the *Rps18* housekeeping gene, in cDNA obtained from N2a neurons transduced with either EGFP (EGFP), E2F4DN-6xHis (E2F4DN) or E2F4CA-6xHis (E2F4CA) treated for the indicated time periods with 10 µM CPT. *p<0.05; **p<0.01 (One-way ANOVA). **B** Percentage of active caspase-3-positive N2a neurons transduced with either EGFP or a *Cited2*-specific shRNA (shCITED2) and cultured for 32 h with 10 µM CPT, related to total DAPI-positive nuclei. **C** Percentage of active caspase-3-positive N2a neurons transduced with either 120 IU/cell of an Ad5.EGFP vector (EGFP), 60 IU/cell of an Ad5.E2F4DN-6xHis vector plus 60 IU/cell of an Ad5.EGFP vector (E2F4DN), 60 IU/cell of an Ad5 vector expressing a *Cited2*-specific shRNA plus 60 IU/cell of an Ad5.EGFP vector (shCITED2), or 60 IU/cell of an Ad5.E2F4DN-6xHis vector plus 60 IU/cell of an Ad5 vector expressing a *Cited2*-specific shRNA (E2F4DN/shCITED2), and cultured for 32 h or 48 h with 10 µM CPT, related to total DAPI-positive nuclei. *p<0.05; **p<0.01; ***p<0.001 (Two-way ANOVA, followed by Tukey’s HSD post-hoc test).

To test the participation of Cited2 in the prevention by E2F4DN of CPT-dependent cell death, we used an adenoviral vector carrying a *Cited2*-specific shRNA (Ad5-m-CITED2-shRNA). The capacity of this vector to co-express EGFP, allowed us to verify that ∼75 % of N2a neurons were transduced (Supplementary Fig. 8a). As expected, this shRNA was able to downregulate the expression of endogenous *cited2* mRNA in N2a neurons (Supplementary Fig. 8b). The transduction of N2a neurons with the Ad5-m-CITED2-shRNA vector confirmed that Cited2 is required for the prevention of CPT-dependent cell death by E2F4DN in these cells. As a first evidence, the presence of the *Cited2*-specific shRNA was able to potentiate the apoptotic effect of CPT (Fig. 7C). Moreover, we verified that Cited2 participates in the antiapoptotic signaling pathway regulated by E2F4DN, as the knockdown of *Cited2* in CPT-treated N2a neurons prevented the prosurvival effect of E2F4DN at both 32 h and 48 h (Fig. 7D).

## Discussion

In this study we have demonstrated that the transcription factor E2F4 can be detected in different phosphorylation states in N2a neurons, showing a spatial pattern that changes when the nucleus is compared to the cytosol. The phosphorylation pattern of exogenous E2F4 changes in response to CPT treatment, due to the activation of p38^MAPK^, a stress kinase that phosphorylates E2F4 in the cytosol within its Thr248/Thr250 motif. This phosphorylation, independently of the phosphorylation state of E2F4 in other residues, reduces the capacity of E2F4 to induce *Cited2* expression, thus favoring CPT-dependent apoptosis. A phosphomimetic form of E2F4 containing the Thr248Glu/Thr250Glu mutation acts synergistically with CPT to induce apoptosis in N2a neurons, although with reduced potency compared to E2F1. In contrast, E2F4DN favors the survival of N2a neurons even under strong CPT-triggered genotoxic stress. This stresses the relevance of the phosphorylation of E2F4 by p38^MAPK^ at the Thr248/Thr250 motif in the apoptotic effect of E2F4 in N2a neurons subjected to genotoxic stress (Fig. 8).

**Fig. 8.**
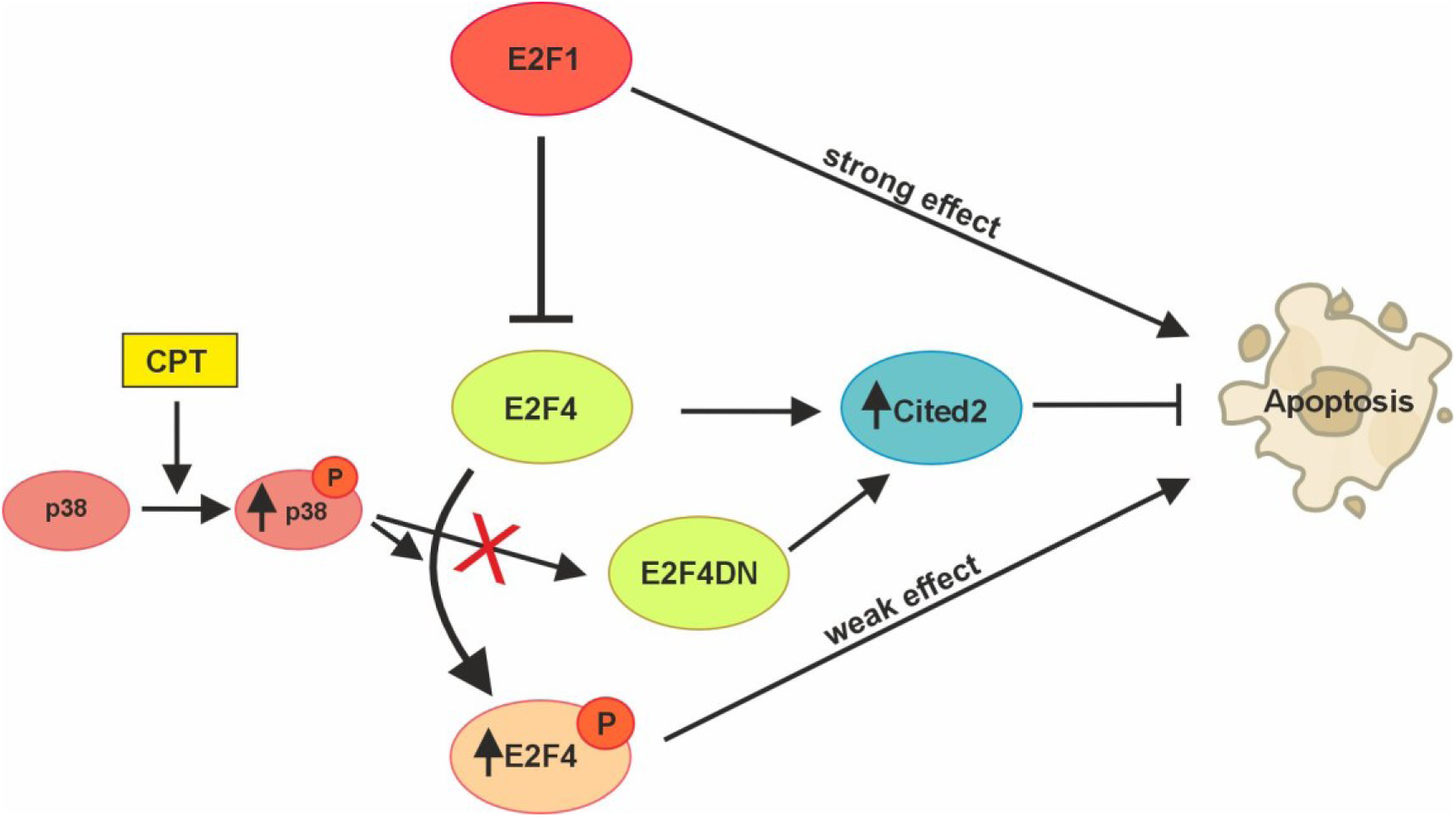
Scheme illustrating the proposed mechanism of E2F4DN to favor the survival of CPT-treated N2a neurons. Under normal conditions, E2F4 facilitates the expression of Cited2, which trigger antiapoptotic signaling in N2a neurons. Genotoxic stress induced by CPT leads to p38^MAPK^ activation, which phosphorylates E2F4 at the Thr249/Thr251 motif. This facilitates a weak proapoptotic effect of E2F4 that mimics the effect of E2F1 as a strong proapoptotic trigger in CPT-treated cells. E2F4DN, which cannot be phosphorylated at the Thr249/Thr251 motif, maintains the capacity of E2F4 to induce Cited2 expression and the antiapoptotic effect of the latter.

We have shown that, in N2a neurons, E2F4 can be detected in different phosphorylation states. This confirms previous results in colorectal adenomas and intestinal epithelial cells [17], in C33A and U937 cells [40], as well as a study in mouse embryonic stem cells showing that E2F4 contains three threonines, one tyrosine, and six serines susceptible to become phosphorylated, in addition to Thr249 [18]. Remarkably, the incubation of the cell extracts with AP resulted in a shift towards two high mobility bands, suggesting that two main forms of E2F4 are present in N2a neurons, likely differing from each other by a covalent modification. This contrasts with the results from [40], who found only one band in C33A cells treated with λ phosphatase. In N2a neurons, E2F4 could be covalently modified by acetylation of its K96 residue [8], a modification that would theoretically result in a higher mobility when resolved in SDS-PAGE gels [62]. Alternatively, the cleavage of E2F4 in N2a neurons, as previously shown for E2F1 [63], could also explain the presence of this high-mobility band. Interestingly, we have provided evidence that this modified form of E2F4, can become phosphorylated by p38^MAPK^ to give rise to lower mobility bands since the AP treatment increases its intensity.

Our study also revealed a different phosphorylation pattern of E2F4 when nuclear and cytosolic extracts were compared, with enhancement of low-mobility bands in the former subcellular compartment, whose particular identity changed depending on the E2F4 variant. Therefore, the translocation of E2F4 to the nucleus seems to be associated with its phosphorylation state, as demonstrated by [17] who shown that ERK1/2-mediated phosphorylation of E2F4 on Ser244 and Ser384 promotes its nuclear localization in proliferating intestinal cells.

CPT treatment resulted in the phosphorylation of the modified form of E2F4WT at Thr248, resulting in the increase of the intensity of band *d* in the cytosol. The conclusion that the phosphorylated residue of E2F4 that becomes phosphorylated in response to CPT is Thr248 was supported by the observation that the intensity of band *d* did not change upon CPT treatment in both E2F4DN- and E2F4CA-expressing N2a neurons. In addition, immunostaining of N2a neurons with an antibody specific for the phosphoThr249 epitope of endogenous E2F4 confirmed that CPT induces the phosphorylation of E2F4 at Thr249.

We provide evidence that the phosphorylation of E2F4 in response to CPT is triggered by p38^MAPK^, a finding consistent with previous studies demonstrating that CPT and Irinotecan, a CPT derivative that also inhibits topoisomerase I, can both induce p38^MAPK^ activity in cancer cells [64,65], thus indicating that p38^MAPK^ can be specifically activated in response to DNA damage [66]. In this regard, we have confirmed that CPT treatment of N2a neurons results in time-dependent activation of p38^MAPK^ but not of JNK, a related stress kinase. The activation of p38^MAPK^ by CPT was paralleled by detection by immunocytochemistry of the phosphoThr249 epitope in N2a neurons, an effect that could be prevented with the p38^MAPK^-specific inhibitor SB203580 but not by JNK inhibition with SP600125. Interestingly, SB203580 is a known inhibitor of the isoforms p38α and p38β [67,68], suggesting that these isoforms are those involved in the phosphorylation of E2E4 in response to cellular stress. These results are consistent with our finding that the domain of mouse E2F4DN where the conserved threonines Thr249 and Thr251 are located functionally interacts with p38α but not with JNK1, as evidenced by cell-free kinase assays. Our results also indicate that SB203580 is not able to reduce the phosphorylation levels of E2F4 at 0-4 h. A possible explanation for this observation is that under basal conditions E2F4 is phosphorylated by p38γ or p38δ, while a switch to the p38α/p38β isoforms may take place at 6-8 h. This view is consistent with the fact that the canonic sequence of human p38^MAPK^ employed to generate the mAb used in this study dramatically differs in the p38γ isoform. Therefore, this antibody could not detect the presence of p38γ in the cell extracts illustrated in Fig. 3A.

p38^MAPK^ activation can act as a switch between prosurvival and proapoptotic signaling depending on the strength of the DNA damage [69–71]. In this regard, highly concentrated CPT treatment represents a strong signal leading to p38^MAPK^-dependent apoptosis [64,71]. The effect of CPT is known to be potentiated by E2F1 in a cell cycle-independent manner [2,24,25]. In line with these observations, we found that E2F1-overexpressing N2a neurons underwent apoptosis 48 h after treatment with 10 μM CPT. E2F4 has been shown to regulate the proapoptotic effect of E2F1 [26], but the role of E2F4 phosphorylation in this paradigm was unknown. We have demonstrated in this study that the CPT-dependent apoptotic response can be potentiated by E2F4, although with reduced capacity compared to E2F1, depending on its Thr248 phosphorylation state. The expression of the Thr248/Thr250 phosphomimetic form of E2F4 (E2F4CA), significantly potentiated the CPT-induced apoptosis in N2a neurons. In contrast, E2F4DN, which lacks the capacity to become phosphorylated by p38^MAPK^ as it contains the Thr248Ala/Thr250Ala mutation, prevented this cell death.

Our results have revealed that the mechanism used by E2F4DN to prevent CPT-dependent cell death in N2a neurons does not rely in its capacity to induce the expression of DNA damage repair genes [8,72]. This lack of repair effect of E2F4DN is likely due to the strong DNA damage triggered by the high concentration of CPT that was used, which resulted in a reduction of nuclei showing punctate γH2AX labeling concomitant with an increase of nuclei with a diffuse γH2AX immunostaining. This diffuse pattern has previously been described in response to strong DNA damage such as that observed in response to UV irradiation [73].

We have shown that the lack of apoptotic effect of E2F4DN in CPT-treated N2a neurons likely depends on Cited2, a transcriptional regulator that may lead to either apoptosis of cell survival depending on the cell context [57,58]. This conclusion is based on two different lines of evidence. We have demonstrated that E2F4DN, but not E2F4CA, upregulates *Cited2* expression in the absence of CPT-dependent p38^MAPK^ activation. Furthermore, *Cited2* knockdown with a specific shRNA potentiated the apoptotic effect of CPT and prevented the antiapoptotic effect of E2F4DN. If Cited2 would be acting independently of E2F4, then the increase of apoptosis observed in N2a neurons transduced with shCITED2 in the presence of CPT would have been reduced by E2F4DN co-expression, but this was not the case (Fig. 7C). Same levels of apoptosis were observed in cells expressing shCITED2 and shCITED2/E2F4DN, indicating that Cited2 acts downstream of E2F4DN. Interestingly, our results in N2a neurons differ from those described by a previous study that found an opposed activity of Cited2 in primary cortical neurons [26]. In this latter paradigm, CPT prevents the capacity of E2F4 to repress the capacity of E2F1 to induce the expression of Cited 2, followed by apoptosis.

In summary, we have uncovered in this study a putative mechanism of action of E2F4DN underlying its multifactorial therapeutic capacity as a homeostasis regulator (Fig. 8). This mechanism, which derives from its resistance to become phosphorylated by p38^MAPK^ under genotoxic stress, may be relevant for AD [8–10], since the affected neurons in this disease have been shown to undergo genotoxic stress [74,75].

## Acknowledgements

The authors thank the Cajal Institute core facilities for their important contributions to this work, particularly the Molecular and Cellular Biology Facility, the Cell Culture Facility and the Scientific Image and Microscopy Facility.

## Data and resource availability

All data generated or analyzed during this study are included in this published article and its supplementary information files. Material are available from the corresponding author on reasonable request

## Author contributions statement

Design of the work: José M. Frade. Acquisition of data: Aina M. Llabrés-Mas, Noelia López-Sánchez, Alberto Garrido-García and Vanesa Cano-Daganzo. Analysis and interpretation of data: Aina M. Llabrés-Mas., Noelia López-Sánchez, and José M. Frade. The first draft of the manuscript was written by José M. Frade and all authors commented on previous versions of the manuscript. All authors read and approved the final manuscript.

## References

1. Wu PY, Lin YC, Chang CL, Lu HT, Chin CH, Hsu TT et al (2009) Functional decreases in P2X7 receptors are associated with retinoic acid-induced neuronal differentiation of Neuro-2a neuroblastoma cells. Cell Signal 21:881–891. 10.1016/j.cellsig.2009.01.036

2. Zhang Y, Song X, Herrup K (2020) Context-dependent functions of E2F1: cell cycle, cell death, and DNA damage repair in cortical neurons. Mol Neurobiol 57:2377–2390. 10.1007/s12035-020-01887-5

3. Conboy CM, Spyrou C, Thorne NP, Wade EJ, Barbosa-Morais NL, Wilson MD, et al (2007) Cell cycle genes are the evolutionarily conserved targets of the E2F4 transcription factor. PLoS One 2:e1061. 10.1371/journal.pone.0001061

4. Lee BK, Bhinge AA, Iyer VR (2011) Wide-ranging functions of E2F4 in transcriptional activation and repression revealed by genome-wide analysis. Nucl Acids Res 39:3558–3573. 10.1093/nar/gkq1313

5. Stevaux O, Dyson NJ (2002) A revised picture of the E2F transcriptional network and RB function. Curr Op Cell Biol 14:684–691

6. Crosby ME, Almasan, A (2004) Opposing roles of E2Fs in cell proliferation and death. Cancer Biol Ther 3:1208–1211

7. Hsu J, Sage J (2016) Novel functions for the transcription factor E2F4 in development and disease. Cell Cycle 15:3183–3190

8. Ramón-Landreau M, Sánchez-Puelles C, López-Sánchez N, Lozano-Ureña A, Llabrés-Mas AM, Frade JM (2022) E2F4DN transgenic mice: a tool for the evaluation of E2F4 as a therapeutic target in neuropathology and brain aging. Int J Mol Sci 23:12093

9. López-Sánchez N, Garrido-García A, Ramón-Landreau M, Cano-Daganzo V, Frade JM (2021) E2F4-based gene therapy mitigates the phenotype of the Alzheimer’s disease mouse model 5xFAD. Neurotherapeutics 18:2484–2503

10. López-Sánchez N, Ramón-Landreau M, Trujillo C, Garrido-García A, Frade JM (2022) A mutant variant of E2F4 triggers multifactorial therapeutic effects in 5xFAD mice. Mol Neurobiol 59:3016–3039

11. Ding J, Kong W, Mou X, Wang S (2019) Construction of Transcriptional Regulatory Network of Alzheimer’s Disease Based on PANDA Algorithm. Interdiscip Sci Comput Life Sci 11:226–236

12. Li P, Marshall L, Oh G, Jakubowski JL, Groot D, He Y, et al (2019) Epigenetic dysregulation of enhancers in neurons is associated with Alzheimer’s disease pathology and cognitive symptoms. Nat Commun 10:2246

13. Caldwell AB, Liu Q, Schroth GP, Galasko DR, Yuan SH, Wagner SL, et al (2020) Dedifferentiation and neuronal repression define familial Alzheimer’s disease. Sci Adv 6:eaba5933. 10.1126/sciadv.aba5933

14. Bottero V, Potashkin JA (2019) Meta-analysis of gene expression changes in the blood of patients with mild cognitive impairment and Alzheimer’s disease dementia. Int J Mol Sci 20:5403

15. Nguyen HD (2022) Combination of donepezil and memantine attenuated cognitive impairment induced by mixed endocrine-disrupting chemicals: an in silico study. Neurotox Res 40:2072–2088. 10.1007/s12640-022-00591-7

16. Morillo SM, Abanto EP, Román MJ, Frade JM (2012) Nerve growth factor-induced cell cycle reentry in newborn neurons is triggered by p38MAPK-dependent E2F4 phosphorylation. Mol Cell Biol 32:2722–2737

17. Paquin MC, Cagnol S, Carrier JC, Leblanc C, Rivard N (2013) ERK-associated changes in E2F4 phosphorylation, localization and transcriptional activity during mitogenic stimulation in human intestinal epithelial crypt cells. BMC Cell Biol 14:33. 10.1186/1471-2121-14-33

18. Hsu J, Arand J, Chaikovsky A, Mooney NA, Demeter J, Brison CM, et al (2019) E2F4 regulates transcriptional activation in mouse embryonic stem cells independently of the RB family. Nat Commun 10:2939. 10.1038/s41467-019-10901-x

19. Oakley H, Cole SL, Logan S, Maus E, Shao P, Craft J, et al (2006) Intraneuronal beta-amyloid aggregates, neurodegeneration, and neuron loss in transgenic mice with five familial Alzheimer’s disease mutations: potential factors in amyloid plaque formation. J Neurosci 26:10129–10140. 10.1523/JNEUROSCI.1202-06.2006

20. Richard BC, Kurdakova A, Baches S, Bayer TA, Weggen S, Wirths O (2015) Gene dosage dependent aggravation of the neurological phenotype in the 5xFAD mouse model of Alzheimer’s disease. J Alzheimers Dis 45:1223–1236. 10.3233/JAD-143120

21. Sánchez-Puelles C, Perea G, Frade JM (2021) E2F4DN-based gene therapy recovers long-term potentiation and hippocampal-dependent memory in homozygous 5xFAD mice. Alzheimers Dement 17(Suppl.9):e057668

22. Mathews RA, Johnson TC, Hudson JE (1976) Synthesis and turnover of plasma-membrane proteins and glycoproteins in a neuroblastoma cell line. Biochem J 154:57–64. 10.1042/bj1540057

23. Morris EJ, Geller HM (1996) Induction of neuronal apoptosis by camptothecin, an inhibitor of DNA topoisomerase-I: evidence for cell cycle-independent toxicity. J Cell Biol 134:757–770. 10.1083/jcb.134.3.757

24. Hofland K, Petersen BO, Falck J, Helin K, Jensen PB, Sehested M (2000) Differential cytotoxic pathways of topoisomerase I and II anticancer agents after overexpression of the E2F-1/DP-1 transcription factor complex. Clin Cancer Res 6:1488–1497

25. Dong YB, Yang HL, McMasters KM (2003) E2F-1 overexpression sensitizes colorectal cancer cells to camptothecin. Cancer Gene Ther 10:168–178. 10.1038/sj.cgt.7700565

26. Huang T, González YR, Qu D, Huang E, Safarpour F, Wang E, et al (2019) The pro-death role of Cited2 in stroke is regulated by E2F1/4 transcription factors. J Biol Chem 294:8617–8629. 10.1074/jbc.RA119.007941

27. Miravet S, Ontiveros M, Piedra J, Penalva C, Monfar M, Chillón M (2014) Construction, production, and purification of recombinant adenovirus vectors. Methods Mol Biol 1089:159–173. 10.1007/978-1-62703-679-5_12

28. Hsiang YH, Hertzberg R, Hecht S, Liu LF (1985) Camptothecin induces protein-linked DNA breaks via mammalian DNA topoisomerase I. J Biol Chem 260:14873–14878

29. Liu LF, Wang JC (1987) Supercoiling of the DNA template during transcription. Proc Natl Acad Sci USA 84:7024–7027. 10.1073/pnas.84.20.7024

30. Cuadrado A, Nebreda AR (2010) Mechanisms and functions of p38 MAPK signalling. Biochem J 429:403–417

31. Bennett BL, Sasaki DT, Murray BW, O’Leary EC, Sakata ST, Xu W, et al (2001) SP600125, an anthrapyrazolone inhibitor of Jun N-terminal kinase. Proc Natl Acad Sci USA 98:13681–13686

32. Kanning KC, Hudson M, Amieux PS, Wiley JC, Bothwell M, Schecterson LC (2003) Proteolytic processing of the p75 neurotrophin receptor and two homologs generates C-terminal fragments with signaling capability. J Neurosci 23:5425–5436. 10.1523/JNEUROSCI.23-13-05425.2003

33. Muñoz JP, Sánchez JR, Maccioni RB (2003) Regulation of p27 in the process of neuroblastoma N2A differentiation. J Cell Biochem 89:539–549. 10.1002/jcb.10525

34. Wu Z, Zheng S, Yu Q (2009) The E2F family and the role of E2F1 in apoptosis. Int J Biochem Cell Biol 41:2389–2397. 10.1016/j.biocel.2009.06.004

35. Tremblay RG, Sikorska M, Sandhu JK, Lanthier P, Ribecco-Lutkiewicz M, Bani-Yaghoub M (2010) Differentiation of mouse Neuro 2A cells into dopamine neurons. J Neurosci Methods 186:60–67. 10.1016/j.jneumeth.2009.11.004

36. Namsi A, Nury T, Hamdouni H, Yammine A, Vejux A, Vervandier-Fasseur D, et al (2018) Induction of neuronal differentiation of murine N2a cells by two polyphenols present in the mediterranean diet mimicking neurotrophins activities: resveratrol and apigenin. Diseases 6:67. 10.3390/diseases6030067

37. Okabe S, Hirokawa N (1989) Rapid turnover of microtubule-associated protein MAP2 in the axon revealed by microinjection of biotinylated MAP2 into cultured neurons. Proc Natl Acad Sci USA 86:4127–4131. 10.1073/pnas.86.11.4127

38. Paonessa F, Latifi S, Scarongella H, Cesca F, Benfenati F (2013) Specificity protein 1 (Sp1)-dependent activation of the synapsin I gene (SYN1) is modulated by RE1-silencing transcription factor (REST) and 5’-cytosine-phosphoguanine (CpG) methylation. J Biol Chem 288:3227–3239. 10.1074/jbc.M112.399782

39. Kügler S, Meyn L, Holzmüller H, Gerhardt E, Isenmann S, Schulz JB, et al (2001) Neuron-specific expression of therapeutic proteins: evaluation of different cellular promoters in recombinant adenoviral vectors. Mol Cell Neurosci 17:78–96

40. Ginsberg D, Vairo G, Chittenden T, Xiao ZX, Xu G, Wydner KL, et al (1994) E2F-4, a new member of the E2F transcription factor family, interacts with p107. Genes Dev 8:2665–2679. 10.1101/gad.8.22.2665

41. Fagan R, Flint KJ, Jones N (1994) Phosphorylation of E2F-1 modulates its interaction with the retinoblastoma gene product and the adenoviral E4 19 kDa protein. Cell 78:799–811. 10.1016/s0092-8674(94)90522-3

42. Ivanova IA, Nakrieko KA, Dagnino L (2009) Phosphorylation by p38 MAP kinase is required for E2F1 degradation and keratinocyte differentiation. Oncogene 28:52–62. 10.1038/onc.2008.354

43. Yang SH, Liu ML, Tien CF, Chou SJ, Chang RY (2009) Glyceraldehyde-3-phosphate dehydrogenase (GAPDH) interaction with 3’ ends of Japanese encephalitis virus RNA and colocalization with the viral NS5 protein. J Biomed Sci 16:40. 10.1186/1423-0127-16-40

44. Zhu X, Rottkamp CA, Boux H, Takeda A, Perry G, Smith MA (2000) Activation of p38 kinase links tau phosphorylation, oxidative stress, and cell cycle-related events in Alzheimer disease. J Neuropathol Exp Neurol 59:880–888. 10.1093/jnen/59.10.880

45. Sun A, Liu M, Nguyen XV, Bing G (2003) P38 MAP kinase is activated at early stages in Alzheimer’s disease brain. Exp Neurol 183:394–405. 10.1016/s0014-4886(03)00180-8

46. Savage MJ, Lin YG, Ciallella JR, Flood DG, Scott RW (2002) Activation of c-Jun N-terminal kinase and p38 in an Alzheimer’s disease model is associated with amyloid deposition. J Neurosci 22:3376–3385. 10.1523/JNEUROSCI.22-09-03376.2002

47. Colié S, Sarroca S, Palenzuela R, Garcia I, Matheu A, Corpas R, et al (2017) Neuronal p38α mediates synaptic and cognitive dysfunction in an Alzheimer’s mouse model by controlling β-amyloid production. Sci Rep 7:45306. 10.1038/srep45306

48. Raingeaud J, Gupta S, Rogers JS, Dickens M, Han J, Ulevitch RJ, et al (1995) Pro-inflammatory cytokines and environmental stress cause p38 mitogen-activated protein kinase activation by dual phosphorylation on tyrosine and threonine. J Biol Chem 270:7420–7426. 10.1074/jbc.270.13.7420

49. Cuenda A, Rouse J, Doza YN, Meier R, Cohen P, Gallagher TF, et al (1995) SB 203580 is a specific inhibitor of a MAP kinase homologue which is stimulated by cellular stresses and interleukin-1. FEBS Lett 364:229–233

50. Gould GW, Cuenda A, Thomson FJ, Cohen P (1995) The activation of distinct mitogen-activated protein kinase cascades is required for the stimulation of 2-deoxyglucose uptake by interleukin-1 and insulin-like growth factor-1 in KB cells. Biochem J 311:735–738

51. Fusaro G, Wang S, Chellappan S (2002) Differential regulation of Rb family proteins and prohibitin during camptothecin-induced apoptosis. Oncogene 21:4539–4548. 10.1038/sj.onc.1205551

52. Park DS, Morris EJ, Bremner R, Keramaris E, Padmanabhan J, Rosenbaum M, et al (2000) Involvement of retinoblastoma family members and E2F/DP complexes in the death of neurons evoked by DNA damage. J Neurosci 20:3104–3114. 10.1523/JNEUROSCI.20-09-03104.2000

53. Stevens C, La Thangue NB (2003) E2F and cell cycle control: a double-edged sword. Arch Biochem Biophys 412:157–169. 10.1016/s0003-9861(03)00054-7

54. Fridman JS, Lowe SW (2003) Control of apoptosis by p53. Oncogene 22:9030–9040. 10.1038/sj.onc.1207116

55. Boon NJ, Oliveira RA, Körner PR, Kochavi A, Mertens S, Malka Y, et al (2024) DNA damage induces p53-independent apoptosis through ribosome stalling. Science 384:785–792. 10.1126/science.adh7950

56. Mah LJ, El-Osta A, Karagiannis TC (2010) γH2AX: a sensitive molecular marker of DNA damage and repair. Leukemia 24:679–686. 10.1038/leu.2010.6

57. Liu YC, Chang PY, Chao CC (2015) CITED2 silencing sensitizes cancer cells to cisplatin by inhibiting p53 trans-activation and chromatin relaxation on the ERCC1 DNA repair gene. Nucleic Acids Res 43:10760–10781. 10.1093/nar/gkv934

58. Wu ZZ, Sun NK, Chao CC (2011) Knockdown of CITED2 using short-hairpin RNA sensitizes cancer cells to cisplatin through stabilization of p53 and enhancement of p53-dependent apoptosis. J Cell Physiol 226:2415–2428

59. Mattes K, Berger G, Geugien M, Vellenga E, Schepers H (2017) CITED2 affects leukemic cell survival by interfering with p53 activation. Cell Death Dis 8:e3132. 10.1038/cddis.2017.548

60. Su D, Song JX, Gao Q, Guan L, Li Q, Shi C, et al (2016) Cited2 participates in cardiomyocyte apoptosis and maternal diabetes-induced congenital heart abnormality. Biochem Biophys Res Commun 479:887–892. 10.1016/j.bbrc.2016.09.101

61. Yoshida T, Sekine T, Aisaki K, Mikami T, Kanno J, Okayasu I (2011) CITED2 is activated in ulcerative colitis and induces p53-dependent apoptosis in response to butyric acid. J Gastroenterol 46:339–349. 10.1007/s00535-010-0355-9

62. Georgieva EI, Sendra R (1999) Mobility of acetylated histones in sodium dodecyl sulfate-polyacrylamide gel electrophoresis. Anal Biochem 269:399–402. 10.1006/abio.1999.4050

63. Zyskind JW, Wang Y, Cho G, Ting JH, Kolson DL, Lynch DR, et al (2015) E2F1 in neurons is cleaved by calpain in an NMDA receptor-dependent manner in a model of HIV-induced neurotoxicity. J Neurochem 132:742–755. 10.1111/jnc.12956.

64. Lee S, Lee HS, Baek M, Lee DY, Bang YJ, Cho HN, et al (2002) MAPK signaling is involved in camptothecin-induced cell death. Mol Cells 14:348–354.

65. Pranteda A, Piastra V, Stramucci L, Fratantonio D, Bossi G (2020) The p38 MAPK Signaling Activation in Colorectal Cancer upon Therapeutic Treatments. Int J Mol Sci 21:2773. 10.3390/ijms21082773

66. Raman M, Earnest S, Zhang K, Zhao Y, Cobb MH (2007) TAO kinases mediate activation of p38 in response to DNA damage. EMBO J 26:2005–2014. 10.1038/sj.emboj.7601668

67. Sabio G, Reuver S, Feijoo C, Hasegawa M, Thomas GM, Centeno F, et al (2004) Stress- and mitogen-induced phosphorylation of the synapse-associated protein SAP90/PSD-95 by activation of SAPK3/p38gamma and ERK1/ERK2. Biochem J 380:19–30. 10.1042/BJ20031628

68. Kuma Y, Sabio G, Bain J, Shpiro N, Márquez R, Cuenda A (2005) BIRB796 inhibits all p38 MAPK isoforms in vitro and in vivo. J Biol Chem 280:19472–19479. 10.1074/jbc.M414221200

69. Phong MS, Van Horn RD, Li S, Tucker-Kellogg G, Surana U, Ye XS (2010) p38 mitogen-activated protein kinase promotes cell survival in response to DNA damage but is not required for the G(2) DNA damage checkpoint in human cancer cells. Mol Cell Biol 30:3816–3826. 10.1128/MCB.00949-09

70. Gong X, Liu A, Ming X, Deng P, Jiang Y (2010) UV-induced interaction between p38 MAPK and p53 serves as a molecular switch in determining cell fate. FEBS Lett 584:4711–4716. 10.1016/j.febslet.2010.10.057

71. Rudolf E, Kralova V, Rudolf K, John S (2013) The role of p38 in irinotecan-induced DNA damage and apoptosis of colon cancer cells. Mutat Res 741–742:27-34. 10.1016/j.mrfmmm.2013.02.002

72. Ren B, Cam H, Takahashi Y, Volkert T, Terragni J, Young RA, et al (2002) E2F integrates cell cycle progression with DNA repair, replication, and G_2_/M checkpoints. Genes Dev 16:245–256. 10.1101/gad.949802

73. Marti TM, Hefner E, Feeney L, Natale V, Cleaver JE. (2006) H2AX phosphorylation within the G1 phase after UV irradiation depends on nucleotide excision repair and not DNA double-strand breaks. Proc Natl Acad Sci USA 103:9891–9896. 10.1073/pnas.0603779103

74. Nelson TJ, Xu Y (2023) Sting and p53 DNA repair pathways are compromised in Alzheimer’s disease. Sci Rep 13:8304. 10.1038/s41598-023-35533-6

75. Kisby GE, Wilson DM 3rd, Spencer PS (2024) Introducing the Role of Genotoxicity in Neurodegenerative Diseases and Neuropsychiatric Disorders. Int J Mol Sci 25:7221. 10.3390/ijms25137221

