## Supplementary Tables and Figures for "Resistance of E2F4DN to p38^MAPK^ phosphorylation attenuates DNA damage-induced neuronal death via Cited2"

### **Supplementary Table 1.** Primary antibodies

| Protein | Manufacturer | Reference | Host | Application | Dilution |
|---|---|---|---|---|---|
| **E2F4** | Millipore | MABE160 | Mouse | ICQ<br>WB | 1:200<br>1:2000 |
| **P-E2F4** | | | Rabbit | ICQ | 1:50 |
| **p-p38** | Cell Signaling | 4511 | Rabbit | WB | 1:1000 |
| **p38** | Cell Signaling | 8690 | Rabbit | WB | 1:2000 |
| **Cleaved Caspase-3 (Asp175)** | Cell Signaling | 9661 | Rabbit | ICQ | 1:150 |
| **6xHis tag** | Abcam | ab18184 | Mouse | WB | 1:1000 |
| **Lamin B1** | Abcam | ab133741 | Rabbit | WB | 1:4000 |
| **Actin** | Millipore | MAB1501R | Mouse | WB | 1:10000 |
| **E2F1** | Santa Cruz | sc-251 | Mouse | WB | 1:500 |
| **p-JNK** | Cell Signaling | 9252 | Rabbit | WB | 1:1000 |
| **JNK** | Cell Signaling | 9251S | Rabbit | WB | 1:1000 |
| **EGFP** | Nacalai Tesque | 04404-84 | Rat | ICQ | 1:500 |
| **γ H2AX** | Abcam | ab26350 | Mouse | ICQ | 1:300 |
| **MAP2** | Abcam | ab5392 | Chicken | ICQ | 1:250 |
| **Synapsin 1** | Abcam | ab64581 | Rabbit | ICQ | 1:400 |
| **Histone H4** | Abcam | ab17036 | Mouse | WB | 1:2000 |
| **GAPDH** | Invitrogen | AM4300 | Mouse | WB | 1:10000 |
| **Sodium channel, pan** | Sigma | S6936 | Rabbit | WB | 1:1000 |

**Supplementary Table 2.** Secondary antibodies

| Antibody | Host | Conjugated with | Manufacturer | Reference | Aplication | Dilution |
|---|---|---|---|---|---|---|
| **Anti-mouse** | Donkey | Alexa 647 | Invitrogen (Life Technologies) | A31571 | ICQ | 1/1000 |
| **Anti-mouse** | Goat | Alexa 594 | Invitrogen (Life Technologies) | A11032 | ICQ | 1/1000 |
| **Anti-mouse** | Goat | Alexa 568 | Invitrogen (Life Technologies) | A21043 | ICQ | 1/1000 |
| **Anti-mouse** | Goat | IRDye 680RD | Li-cor | 926-68070 | WB | 1/7500 |
| **Anti-rabbit** | Goat | Alexa 568 | Invitrogen (Life Technologies) | A11036 | ICQ | 1/1000 |
| **Anti-rabbit** | Donkey | Alexa 488 | Invitrogen (Life Technologies) | A21206 | ICQ | 1/1000 |
| **Anti-rabbit** | Goat | IRDye 800CW | Li-cor | 926-32211 | WB | 1/7500 |
| **Anti-rat** | Goat | Alexa 647 | Invitrogen (Life Technologies) | A21247 | ICQ | 1/1000 |

**Supplementary Table 3.** Oligonucleotides used in qPCR

| Transcript | Species | Position (bp) | NCBI accesión # | Size (bp) | Sequence 5' –> 3' |
|---|---|---|---|---|---|
| ***Rps18*** | *M. musculus* | 120-139 | NM_011296 | 73 | **U:** AATAGCCTTCGCCATCACTG |
| | | 173-192 | | | **D:** GTCTGCTTTCCTCAACACCA |
| ***Cited2*** | *M. musculus* | 878-897 | NM_010828 | 76 | **U:** CGCCCAATGTCATAGACACT |
| | | 932-953 | | | **D:** CGGTCCAAACCCATTTCTATCA |
| ***Syn1*** | *M. musculus* | 2108-2129 | NM_013680.4 | 103 | **U:** AGCTCAACAAATCCCAGTCTCT |
| | | 2191-2210 | | | **D:** CGGATGGTCTCAGCTTTCAC |

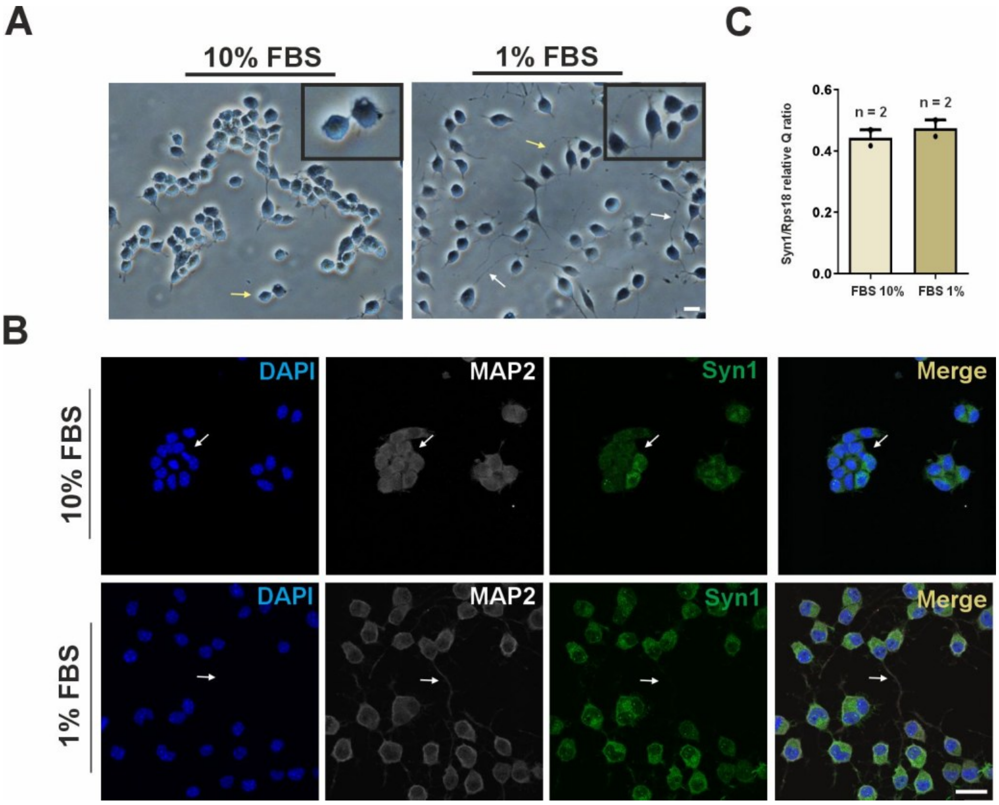


**Supplementary Fig. 1** Differentiation of N2a neuroblastoma cells into N2a neurons. **A** Representative bright field images of N2a neuroblastoma cells cultured with DMEM supplemented with FBS 10% (10 % FBS) or FBS 1% (1% FBS). Box: high magnification to visualize the neurites (arrows) that can be observed in N2a neuroblastoma cells cultured with 1% FBS (i.e. N2a neurons). Bar: 25 μm. **B** Representative immunostaining of N2a neuroblastoma cells cultured with 10% FBS or 1% FBS, using antibodies for the neuronal markers MAP2 (MAP2) and Synapsin 1 (Syn1). Cell nuclei were labeled with DAPI (blue). Arrow: MAP2-positive process. Bar: 25 μm. **C** Quantification of *Syn1* mRNA by real-time quantitative PCR (RT-qPCR) normalized to the *Rps18* housekeeping gene in cDNA obtained from N2a neurons cultured with either FBS 10% or FBS 1%.

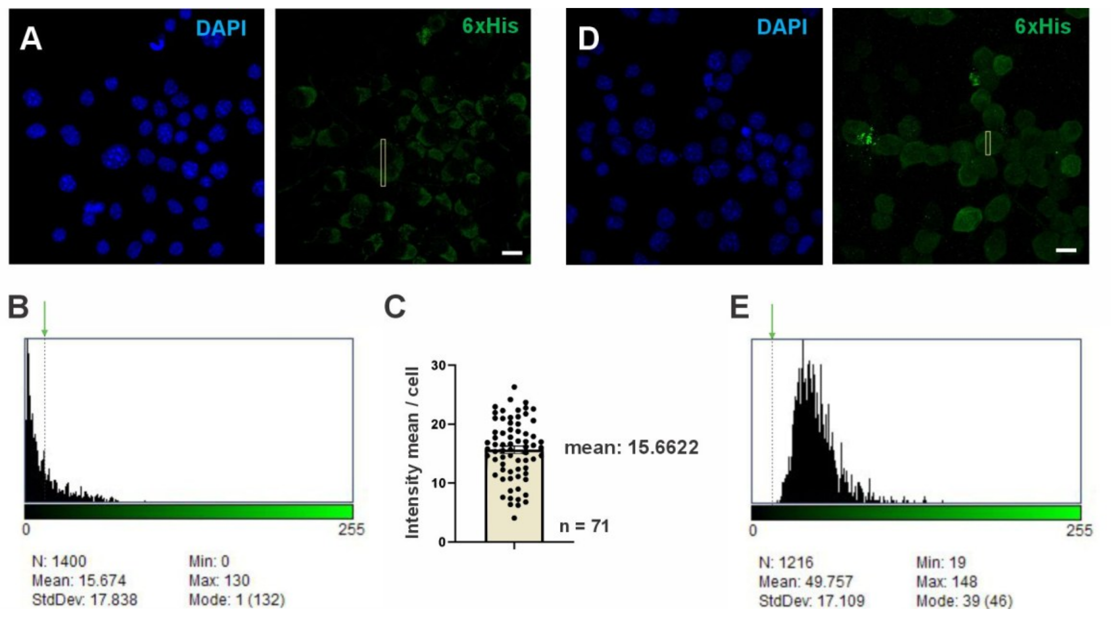


**Supplementary Fig. 2** Method for quantifying Ad5.E2F4WT-6xHis-, Ad5.E2F4DN-6xHis-, and Ad5.E2F4CA-6xHis-transduced cells. **A** Representative image showing 6xHis-specific immunostaining of non-transduced cells. White rectangle: area selected to analyze green fluorescence per cell. **B** Representative histogram of fluorescence intensity per cell. **C** Quantification of mean fluorescence intensity per cell in 71 randomly chosen cells. Average mean intensity per cell obtained: 15.6622 (identified in panels B and E with a dotted line and a green arrow). Transduced cells are considered positive for the 6xHis tag when their mean fluorescence value is above this value. **D** Representative image showing 6xHis-specific immunostaining of Ad5.E2F4WT-6xHis-transduced cells. White rectangle: area selected to analyze green fluorescence per cell. **E** Representative histogram of an Ad5.E2F4WT-6xHis-transduced cell positive for 6xHis tag.

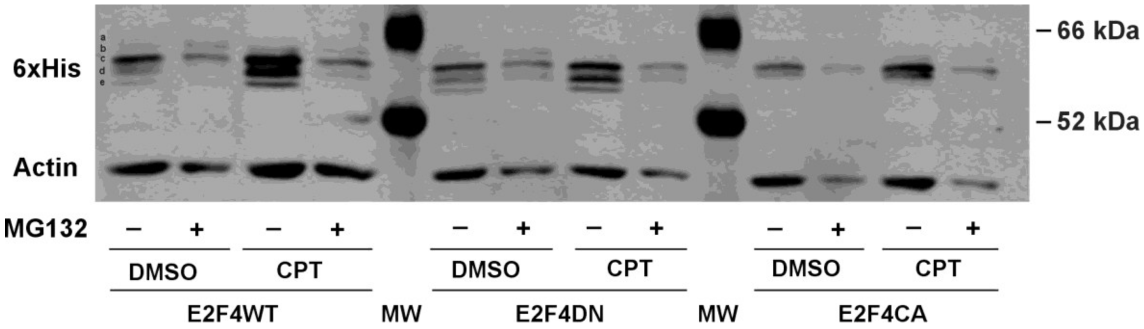


**Supplementary Fig. 3** Proteasome inhibition in E2F4WT, E2F4DN and E2F4Ca-transduced N2A neurons. Representative western blots of the multiband pattern (bands *a*-*e*) in the indicated homogenates from E2F4WT-6xHis- (E2F4WT), E2F4DN-6xHis- (E2F4DN) or E2F4CA-6xHis-transduced N2a neurons (E2F4CA) treated with MG132 and then, 1 h later, treated with CPT for 8 h or maintained under control conditions, as revealed with an anti-6xHis antibody. Actin was used as a loading marker.

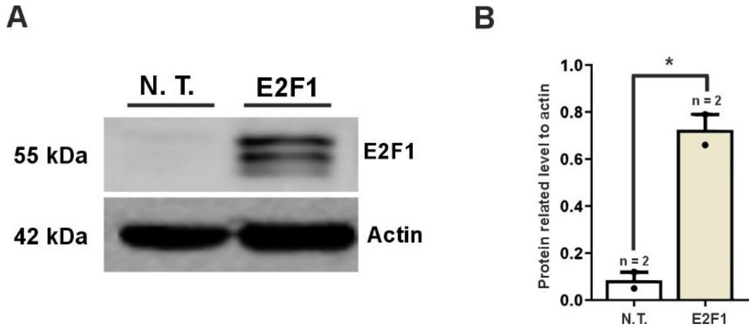


**Supplementary Fig. 4** Overexpression and phosphorylation of E2F1 in N2a neurons. **A** Representative western blot of cell extracts from N2a neurons either non-transduced (N.T.) or transduced with an Ad5.HA-E2F1 vector (E2F1) using antibodies against E2F1 and actin. Notice that exogenous E2F1show a multiple band pattern, likely due to phosphorylation. **B** Quantification of the E2F1 protein level relativized to actin in cell extracts from N2a neurons either non-transduced (N.T.) or transduced with an Ad5.HA-E2F1 vector (E2F1). *p<0.05 (unpaired Student's *t* test).

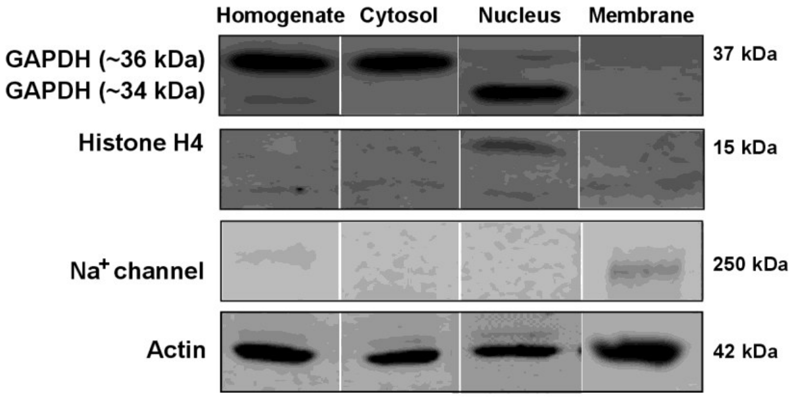


**Supplementary Fig. 5** Cellular fractionation markers in N2a neurons. Gels were overrun to resolve the E2F4 phosphorylation bands (60-70 kDa), thus resulting in the loss of the Histone H4 band due to its extreme high mobility (15 kDa). Therefore, different gels were used to obtain the E2F4 bands and histone localization controls. Representative western blots of whole cell extracts (homogenates) and the indicated subcellular fractions from N2a neurons, using antibodies against GAPDH, Histone H4, sodium channel ($Na^+$ channel), and Actin are shown. Notice that GAPDH can be detected in the cytosolic fraction as a band of ~36 kDa, as previously described [43], while in the nucleus it can be observed as a prominent band of higher mobility (~34 kDa) and a slight band of identical mobility to cytosolic GAPDH (~36 kDa). Histone H4 is specific for the nuclear fraction, and sodium channel only appears in the membrane fraction. Actin was used as a loading marker.

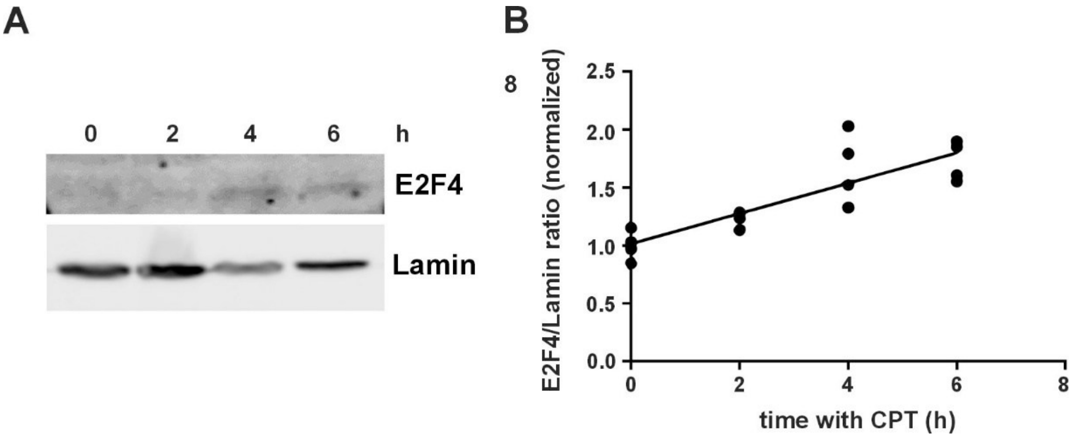


**Supplementary Fig. 6** Increase of E2F4 protein in CPT-treated N2a neurons. **A** Representative western blot of cell extracts from N2a neurons at the indicated time points after 10 μM CPT treatment, using an anti-E2F4-specific antibody. Lamin B1 (Lamin) was used as a loading marker. **B** Regression analysis of the E2F4 protein levels at the indicated time points after 10 μM CPT treatment. $R^2$=0.9646; p=0.0354.

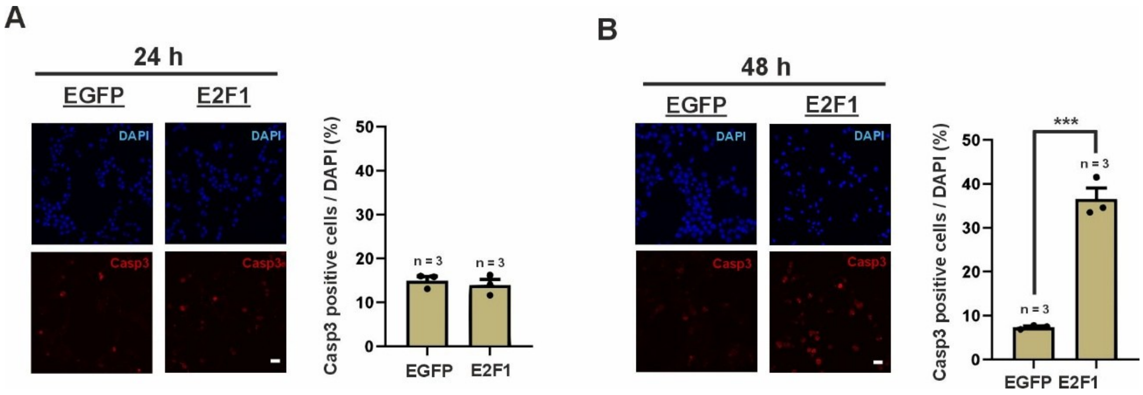


**Supplementary Fig. 7** Regulation of apoptosis by E2F1 in CPT-treated N2a neurons. Percentage of active caspase-3-positive N2a neurons related to total DAPI-positive nuclei at the indicated time points. ***$p<0.005$ (Two-way ANOVA, followed by Tukey's HSD post-hoc test).

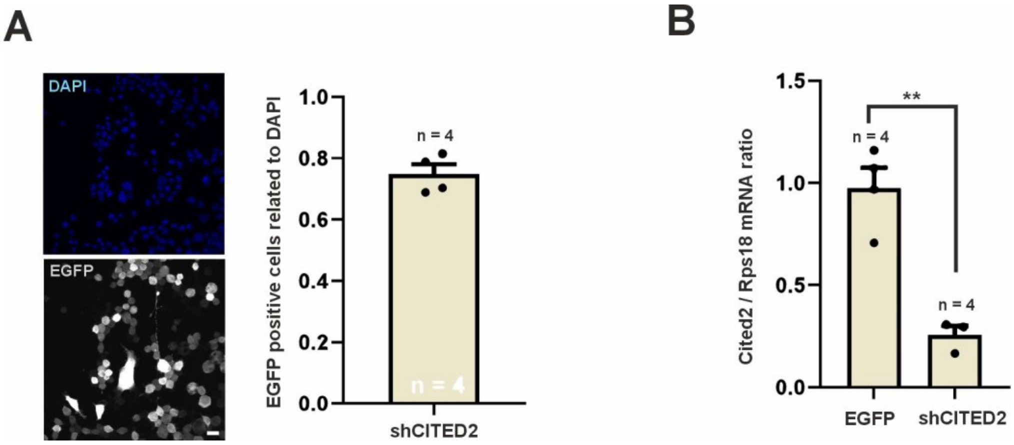


**Supplementary Fig. 8** Regulation of *Cited2* expression by a *Cited2*-specific shRNA in N2a neurons. **A** Left panel: representative immunostaining of N2a neurons transduced with an Ad5 vector co-expressing a *Cited2*-specific shRNA together with EGFP, using antibodies against EGFP. Cell nuclei were labeled with DAPI (blue). Right panel: quantification of EGFP-positive N2a neurons related to DAPI-positive nuclei. Bar: 25 μm. **B** Quantification of *Cited2* mRNA by RT-qPCR normalized to the *Rps18* housekeeping gene in cDNA obtained from N2a neurons transduced with either EGFP or an Ad5 vector co-expressing a *Cited2*-specific shRNA together with EGFP. **p<0.01 (unpaired Student's *t* test).
